# Integrative phosphoproteomics defines two biologically distinct groups of KMT2A rearranged acute myeloid leukaemia with different drug response phenotypes

**DOI:** 10.1101/2022.11.10.515949

**Authors:** Pedro Casado, Ana Rio-Machin, Juho J. Miettinen, Findlay Bewicke-Copley, Kevin Rouault-Pierre, Szilvia Krizsan, Alun Parsons, Vinothini Rajeeve, Farideh Miraki-Moud, David C Taussig, Csaba Bödör, John Gribben, Caroline Heckman, Jude Fitzgibbon, Pedro R. Cutillas

## Abstract

Acute myeloid leukaemia (AML) patients harbouring certain chromosome abnormalities have particularly adverse prognosis. For these patients, targeted therapies have not yet made a significant clinical impact. To understand the molecular landscape of poor prognosis AML we profiled 74 patients from two different centres (in UK and Finland) at the proteomic, phosphoproteomic and drug response phenotypic levels. These data were complemented with transcriptomics analysis for 39 cases. Data integration highlighted a phosphoproteomics signature that define two biologically distinct groups of KMT2A rearranged leukaemia, which we term MLLGA and MLLGB. MLLGA presented increased DOT1L phosphorylation, HOXA gene expression, CDK1 activity and phosphorylation of proteins involved in RNA metabolism, replication and DNA damage when compared to MLLGB and no KMT2A rearranged samples. MLLGA was particularly sensitive to 15 compounds including genotoxic drugs and inhibitors of mitotic kinases and inosine-5-monosphosphate dehydrogenase (IMPDH) relative to other cases. Intermediate-risk KMT2A-MLLT3 cases were mainly represented in a third group closer to MLLGA than to MLLGB. The expression of IMPDH2 and multiple nucleolar proteins was higher in MLLGA and correlated with the response to IMPDH inhibition in KMT2A rearranged leukaemia, suggesting a role of the nucleolar activity in sensitivity to treatment. In summary, our multilayer molecular profiling of AML with poor prognosis and KMT2A-MLLT3 karyotypes identified a phosphoproteomics signature that defines two biologically and phenotypically distinct groups of KMT2A rearranged leukaemia. These data provide a rationale for the potential development of specific therapies for AML patients characterised by the MLLGA phosphoproteomics signature identified in this study.

## INTRODUCTION

Acute Myeloid Leukaemia (AML) is a highly heterogeneous malignancy characterised by impairment of myeloid progenitor cell differentiation, leading to their clonal expansion and, ultimately, bone marrow failure^1^. Despite advances in the development of targeted therapies for AML, current treatments are not curative for most patients^2^. Cases presenting complex karyotypes, KMT2A-rearrangements (excluding KMT2A-MLLT3) and alterations on certain chromosomes (e.g. deletions in chromosomes 5, 7 and 19) and genes (e.g. *TP53* mutations) present particularly poor prognosis and, for these patients, targeted therapies are not yet available^3^. The short overall survival of AML cases with adverse genetic alterations highlights the need of new therapies for these patients^4–7^.

In this work, we subjected a cohort of 74 AML patients with poor prognosis or KMT2A-MLLT3 karyotypes from two different centres (in UK and Finland) to proteomic and phosphoproteomic analysis as well as drug response profiling to 627 drugs. For most patients, sufficient material was also available for additional analysis at the genomic and transcriptomics levels. Proteomics data was additionally mined for proteomic methylation and acetylation post-translational modification analysis. We focused our investigation in the characterization of KMT2A\MLL rearranged AML (KMT2Ar-AML) phosphoproteomics because cells with this karyotype showed distinct phosphoproteomes relative to other poor-risk AML cases.

The *KMT2A* gene, also known as mixed lineage leukaemia or MLL, is located on chromosome 11q23 and encodes a methyltransferase for histone H3 at K4. Balanced chromosome rearrangements between 11q23 and other chromosomes, present in ~5% of de novo AML, generate several distinct KMT2A fusion proteins^8^, the most frequent of which are MLLT3 (not considered an adverse karyotype), MLLT1, MLLT10, ELL and MLLT4^9^. In KMT2Ar-AML, the c-terminal portion of KMT2A – responsible for the methyl transferase activity of the enzyme – is replaced by a region of the fusion partner that leads to the recruitment, probably indirectly in most of the cases, of the DOT1L and TEFb complexes to the KMT2A fusion binding sites. These complexes have Histone H3 K79 methyltransferase and kinase activities, respectively. Aberrantly activated DOT1L and TEFb at regulatory regions of *HOXA* and other KMT2A target genes subvert the transcriptional programmes that promote normal leukaemogenesis^10–12^.

Although the molecular biology of KMT2Ar-AML is relatively well understood, it is not clear whether the distinct KMT2A fusion partners confer cells with different phenotypes. Furthermore, current knowledge has not yet translated into KMT2Ar-AML specific therapies. Targeted personalised therapies require the stratification of cases into phenotypically homogeneous patient groups predicted to respond or not to given treatments. Traditionally, the application of genomic approaches to derive molecular profiles of healthy and pathogenic tissues have driven the development of biomarkers used for patient stratification^13,14^. However, the realisation that non-genomic mechanisms contribute to drug resistance to anti-cancer therapies^15^ has spurred the application of proteomics and phosphoproteomics approaches to rationalise drug response phenotypes^16,17^. These other omic analytical platforms measure the consequences of several layers of regulation (protein expression, modification and direct enzymatic activity) that determine phenotypes^18–21^, and therefore, have the potential to define biomarkers for the stratification of patients into phenotypically homogeneous groups with greater precision than when just using genetic approaches by themselves^22–24^.

Here, we identified a phosphoproteomics signature that defined two biologically distinct groups of KMT2Ar-AML patients with differential sensitivity to multiple approved and experimental drugs. Focusing on the mode of action of the IMPDH inhibitor AVN-944, we found that the sensitivity to this compound was associated to the expression of IMPDH2 in KMT2Ar-AML as well as to the expression of multiple proteins linked to nucleolar biology and RNA metabolism. To determine causality, we stablished that IMPDH inhibition increased the phosphorylation of nucleolar proteins with roles in ribosomal RNA (rRNA) metabolism only in cells sensitive to the inhibitor of this enzyme. Overall, our study uncovers subgroups of biochemically distinct KMT2Ar-AML with differential drug response phenotypes.

## RESULTS

### Overview of multi-omic analysis of poor-risk and KMT2A-MLLT3 AML

Our study initially included 74 patient samples with poor-risk or KMT2A-MLLT3 karyotypes and 4 samples derived from healthy donors (Figure 1a). Samples were collected by the Barts Cancer Institute tissue bank in UK (n=56) and the Institute for Molecular Medicine in Finland (n=18). To increase the robustness of our analysis, we performed a quality control analysis which led to the exclusion of 19 samples from our phospho(proteomic) dataset. Excluded samples were mislabelled as AML (n=2) or presented high number of red blood cells (n=1), T-cells (n=12) or low viability (n=5). In the remaining 55 patient samples (BCI, n=40; FIMM, n=15), the most represented karyotypes were complex karyotype followed by KMT2Ar and alterations in chromosome 7(−7/del(7)) (Figure 1b). In general, KMT2Ar-AML cases were younger than patients in other karyotype groups (Supplementary Figure 1b) and, as previously described^25^, presented leukaemia with morphological features associated to myelomonocytic (M4) or monocytic (M5) differentiation (Supplementary Figure 1b).

**Figure 1.**
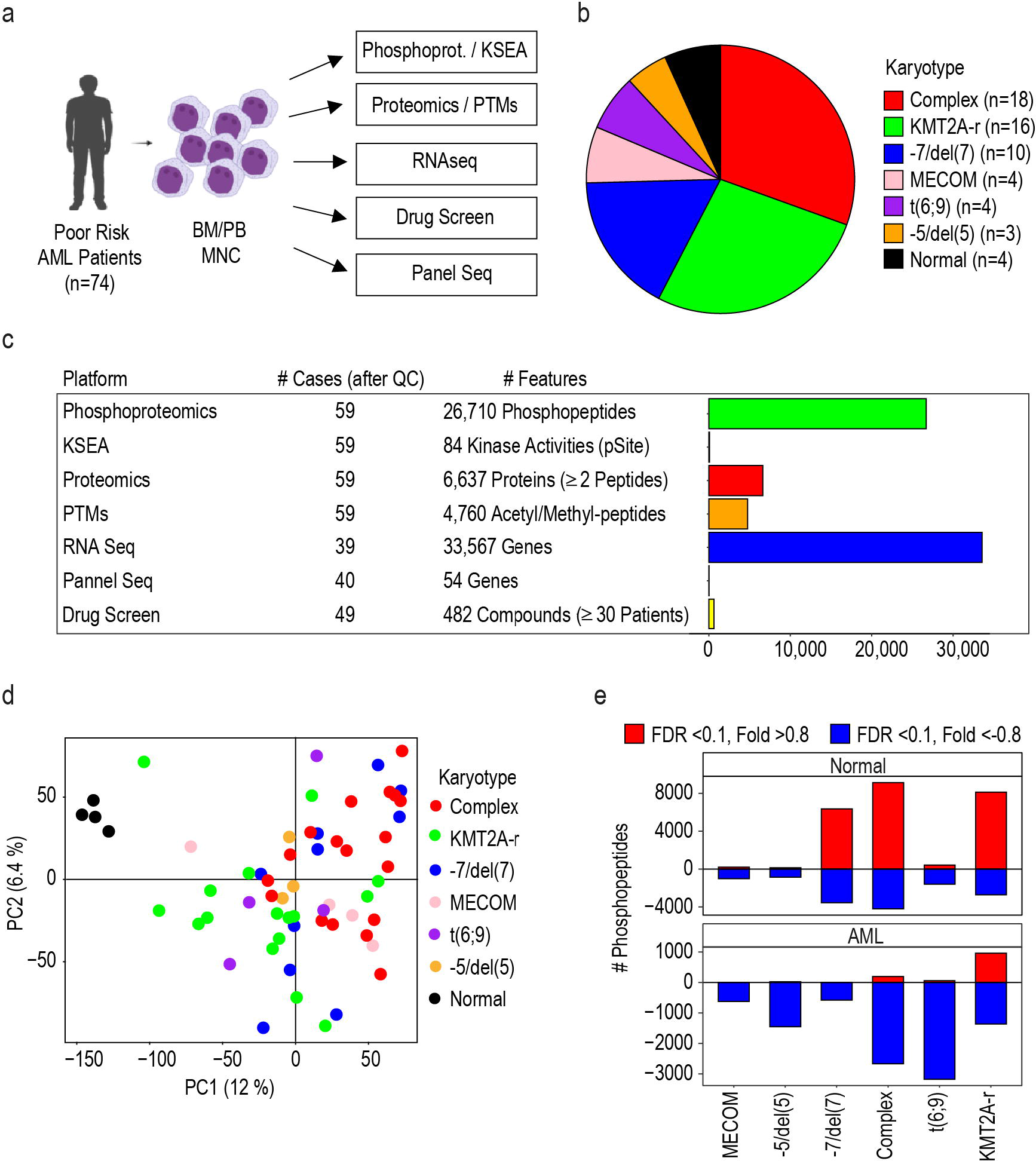
Deep multiomic analysis of poor-risk and KMT2A-MLLT3 AML. **a** Workflow used for the molecular profiling of the patients. **b** Patient cohort as a function of karyotype. **c** Overview of analysed features across the omics layers of the study. **d** Phosphoproteomics separate healthy donor from AML samples in PCA. **e** Number of differentially phosphorylated peptides across karyotype groups and healthy cells. Phosphopeptides were counted when p-value <0.05 and fold change (log2) > 0.8 (positive values) or <-0.8 (negative values). Statistical significance was calculated using unpaired two-sided Student’s t-test. Section **a** was created with BioRender.com.

Mononuclear cells from AML patients and healthy donors were collected from peripheral blood or bone marrow aspirates and subjected to genomics, transcriptomics, proteomics, phosphoproteomics and a PTMs analysis that included peptide methylation and acetylation. Kinase activity was estimated from phosphoproteomics data and cells were subjected to an *ex vivo* drug screening (Figure 1a). Our study quantified 26,710 phosphopeptides 4,760 acetylated or methylated peptides, 6,637 proteins and 84 kinase activities in all 59 individuals (55 patients and 4 healthy donors) (Figure 1c). A total of 33,561 mRNAs were quantified in 39 patients, and 54 frequently mutated genes in AML were sequenced in 40 patients (Figure 1c). Samples from 47 patients and 2 independent healthy donors (PBMCs) were subjected to an *ex vivo* drug screen consisting of 627 compounds assessed across a variable number of patients, with 482 compounds assessed in 30 patients or more (Figure 1c and Supplementary Figure 1c). These results provide the community with the most comprehensive multi-omics dataset of poor-risk AML to date.

### Phosphoproteomics defines two subgroups of KMT2Ar-AML

Targeted therapies require the stratification of patients into phenotypically homogeneous groups. Since phosphorylation regulates the activity of proteins that determine the cell phenotype, we investigated the use of phosphoproteomics as a means to classify poor-risk AML patients. We completed a principal component analysis (PCA) with all identified phosphopeptides to assess the global quality of our phosphoproteomics data. We observed no separation of samples as a function of source (peripheral blood or bone marrow) or origin (UK or Finland) (Supplementary Figure 2a and b), showing that the data was normalized for potential batch effects. In addition, this analysis separated healthy donors from AML samples (Figure 1d). Further characterization of the phosphoproteome of our patient cohort revealed that cells from healthy donors showed a substantial alteration of phosphorylation sites when compared with cells with −7/del(7), complex and KMT2Ar karyotypes (Figure 1e, upper panel). In addition, KMT2Ar-AML cases showed the greater number of significantly increased phosphorylation sites relative to other karyotypes (Figure 1e, lower panel), thus suggesting that the phosphoproteome of KMT2Ar-AML cases is distinct from those of other poor-risk AML cases.

To investigate the nature of the KMT2Ar specific phosphoproteome in more detail, we generated a KMT2Ar signature which we used in a machine learning (ML) approach, based on random forest, to classify AML samples as follows. First, we split our cohort of patients from the BCI into training (n=36) and validation sets (n=8) and used the FIMM cases as an additional verification set (n=15) (Figure 2a and Supplementary Figure 3a). A feature selection process was subsequently applied by comparing the phosphoproteome of cases in the training set with KMT2Ar against samples with other karyotypes (by t-test, Supplementary Figure 3b), from which phosphopeptides with the lowest p-values were selected. To avoid overfitting, we selected a number of features equal to half the number of samples in the training set (n=18) (Figure 2a). To define classes, we then analysed by PCA the resulting KMT2Ar phosphoproteomics signature in all samples in our patient cohort. PC1 separated KMT2Ar-AML from other AML samples (Figure 2b). Sample dispersion in the PCA plot, followed by clustering analysis, indicated that the group with KMT2Ar could be further subdivided into two distinctive groups with two samples separated from the other eight (Figure 2c). We used the cluster of eight samples to define an area in the PCA and we labelled all samples in that area as KMT2A group A (MLLGA) and all samples out of that area as No-MLLGA (Figure 2b). Next, a new random forest classification model, trained using the training set and the phosphoproteomics signature with MLLGA or No-MLLGA as labels, was used to reclassify samples (Figure 2d). Cases that were in the MLLGA area (T5876 and SF13) or did not present KMT2Ar were correctly classified as MLLGA or No-MLLGA, respectively. Four samples in the validation or verification set presented KMT2Ar and were out of the MLLGA. Cases T2353, SF02 and SF16 were clearly separated from the MLLGA area and were classified as No-MLLGA, while sample SF10 that was closed to the MLLGA area was classified as MLLGA and considered as MLLGA in further analyses (Figure 2b and 2d, lower panel). Of note, samples classified as MLLGA (n=11) mainly presented KMT2A fusion proteins involving MLLT4 (n=6) or MLLT10 (n=3), while MLLT3 (n=1) and TET1 (n=1) were the other KMT2A partners represented in this group. Interestingly, analysis of feature importance showed that the most relevant features for the ML model were phosphorylation sites on DOT1L (Figure 2e), a protein highly associated to the biology of KMT2Ar-AML^26^. Together, this analysis uncovered an 18-phosphopeptide signature that separates KMT2Ar-AML cases into two groups based on a ML model in which DOT1L phosphorylation is an important contributor.

**Figure 2.**
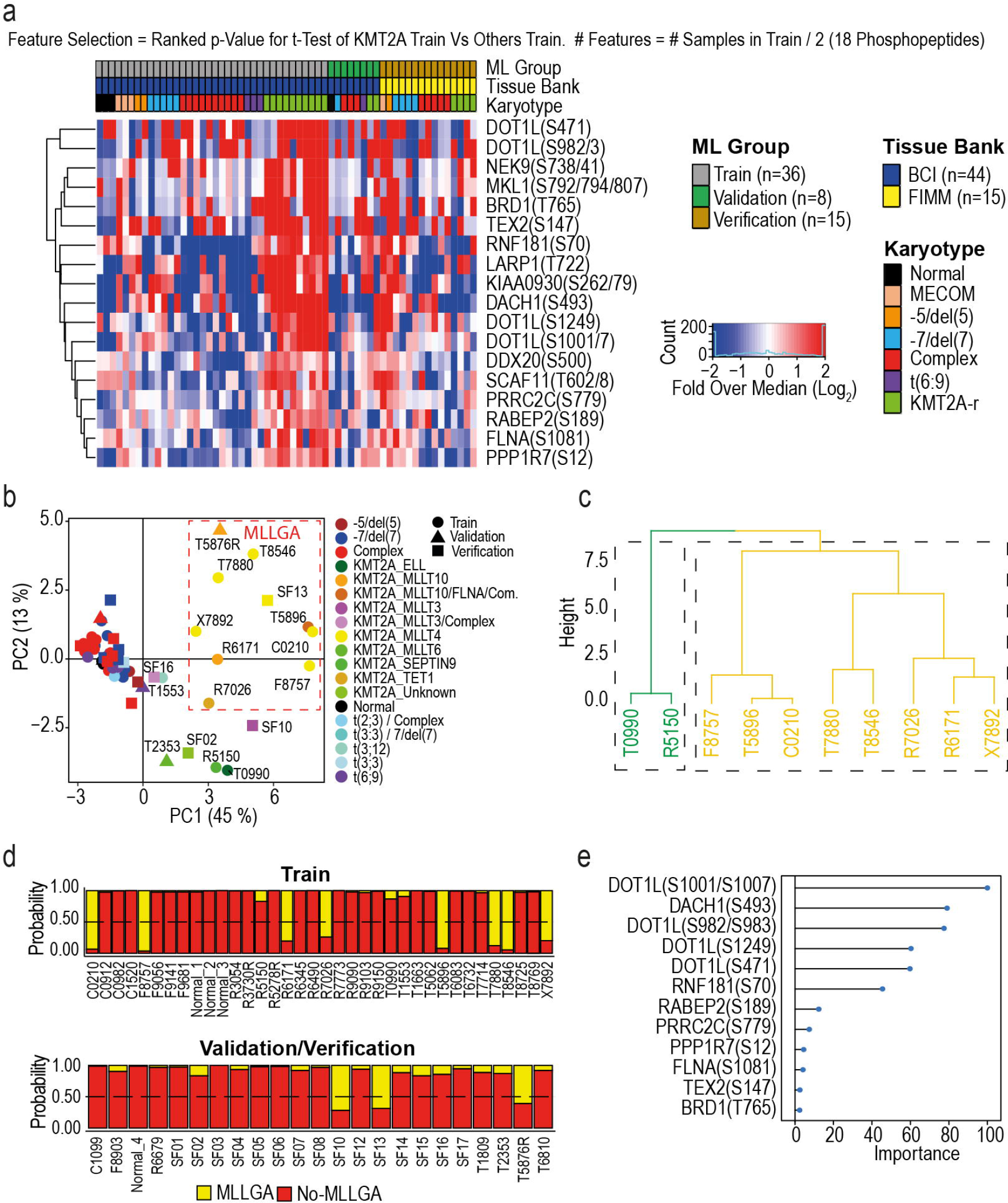
Identification of a phosphoproteomics signature that stratifies KMT2A rearranged leukaemia into two biochemically distinct groups. **a** Phosphoproteomics signature across samples in the training (BCI, UK) and testing (FIMN, Finland) sets. **b** PCA using the phosphoproteomics signature shown in **a**. **c** Definition of the KMT2Ar group MLLGA using hierarchical cluster analysis of PC1 and PC2 (in **b)** in the training set samples. **d** Probability of training and testing set samples to belong to MLLGA or No-MLLGA calculated with a random forest classification model. **e** Rank of feature relevance in the classification model in **d**.

### Distinctive DOT1L-TEFb complex phosphorylation and HOXA gene expression in KMT2Ar groups

Most of the characterised KMT2A fusion proteins directly or indirectly recruit the DOT1L and TEFb complexes to the regulatory regions of target genes^27^ (Figure 3a). Since the phosphorylation of DOT1L was an important contributor to MLLGA in our ML model, we next considered if MLLGA samples presented a specific phosphorylation pattern in DOT1L and TEFb complex components. We found that MLLGA samples presented a significantly increased phosphorylation of sites in DOT1L, MLLT10, MLLT4 and EAF2 and a significantly reduced phosphorylation of AFF4 sites when compared to No-MLL (i.e., non-KMT2Ar cases) and MLLGB (i.e., KMT2Ar-AML cases that were not classified as MLLGA) (Figure 3b). In addition, compared to the No-MLL group, MLLGA and MLLGB presented a significantly reduced phosphorylation of KMT2A at sites located in the c-terminal region that is replaced by the fusion partner (Figure 3b). There was no difference in KMT2A expression between the different groups and protein and phosphorylation site abundances were not significantly correlated (Supplementary Figure 4a and b). Similarly, changes in the protein or mRNA levels of DOT1L and TEFb complex components were not correlated with their extent of phosphorylation (Supplementary Figures 4c and d).

**Figure 3.**
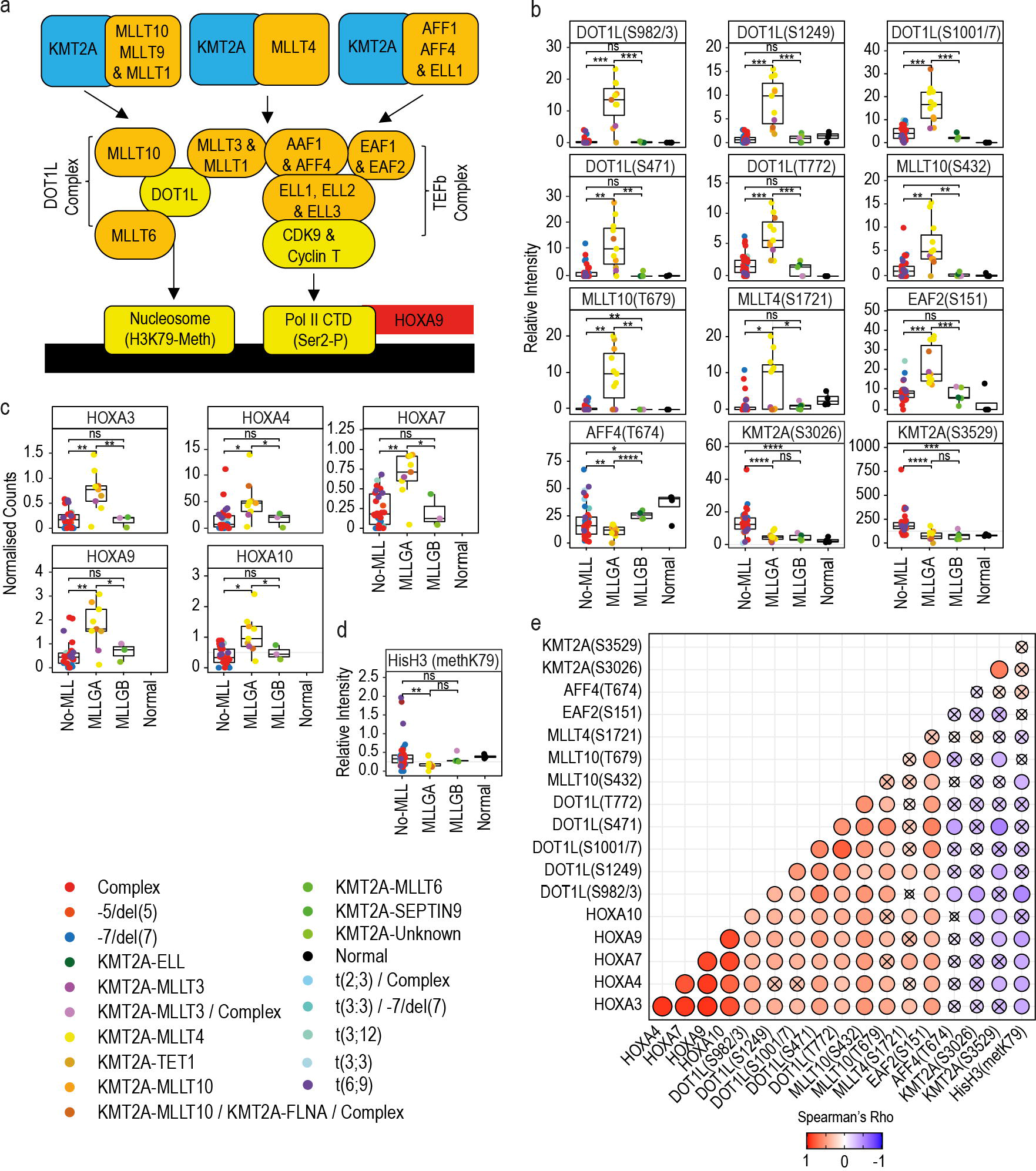
MLLGA increased the phosphorylation of DOT1L and TEFb complex components and the expression of HOXA genes. **a** Schematic of mechanism used by KMT2A fusion proteins to induce HOXA gene expression. **b, c, d** Phosphorylation of DOT1L, TEFb complex components and KMT2A (**b**), HOXA gene expression (**c**) and histone H3 K79 methylation (**d**) across newly identified KMT2Ar groups. **e** Spearman correlation matrix for variables shown in **a** to **d**. **f** HOXA gene expression in dataset obtained from^30^. Data points represent individual patient observations. Boxplots indicate median, 1^st^ and 3^rd^ quartiles. Whiskers extends from the hinge to the largest and lowest value no further than 1.5 times the distance between the 1st and 3rd quartiles (**a**-**d**). For phosphoproteomics and methylation analyses, Normal (n=4), MLLGA (n=11), MLLGB (n=5) and No-MLL (n=39) (**b** and **d**); for mRNA analysis, Normal (n=0), MLLGA (n=9), MLLGB (n=3) and No-MLL (n=27) (**c**). Statistical significance was calculated using unpaired two-sided Student’s t-test. **** p ≤ 0.0001, *** p ≤ 0.001, ** p ≤ 0.01 and * p ≤ 0.05. (**a-d**). Crossed dots indicate no statistically significant correlation (**e**).

DOT1L is an epigenetic modulator that specifically methylates histone H3 at K79^28^. In KMT2Ar-AML, the aberrant activity of DOT1L and TEFb complexes lead to an increase in the expression of genes in the HOXA cluster^11^ (Figure 3a). Consistent with this, we found that that MLLGA significantly increased mRNAs levels for multiple HOXA genes (Figure 3c) and long non-coding RNAs located in the HOXA cluster when compared to MLLGB or No-MLL samples (Supplementary Figure 4e). MLLGA also presented a significantly increased protein expression and phosphorylation of HOXA10 (Supplementary Figure 4f and g). Of note, our data show that MLLGA but not MLLGB presented a reduced methylation of histone H3 at K79 when compared with No-MLL (Figure 3d). We further found that the mRNA levels of multiple HOXA genes positively correlated with the phosphorylation of DOT1L and other components of the DOT1L complex and were inversely correlated with the global methylation of histone H3 at K79 (Figure 3e). Overall, our data on DOT1L and TEFb complex phosphorylation, global H3 methylation at K79 and HOXA gene expression indicate that MLLGA cases have high activity of DOT1L and TEFb at KMT2A target genes.

AML cases with t(9:11) rearrangements generate a KMT2A-MLLT3 fusion protein but these cases are not classified as poor-risk. Intriguingly, however, KMT2A-MLLT3 proteins are also able to recruit the DOT1L and TEFb complexes to the KMT2A target genes^29^. To investigate if KMT2A-MLLT3 cases resemble MLLGA or MLLGB, we generated a new phosphoproteomics dataset for KMT2Ar-AML samples (n=33) that comprised all samples previously classified as MLLGA and MLLGB (n=16). In addition, we analysed new cases containing the following karyotypes: KMT2A-MLLT3 (n=10), MLLT10 (n=1), MLLT4 (n=2), MLLT11 (n=2), t(11;19) rearrangements that generates fusions with ELL or MLLT1 (n=1) and rearrangements involving 11q23 (Unknown; n=1). Although we reduced 60% the amount of starting material due to sample availability limitations, we identified and quantified 10,503 phosphopeptides in this experiment. Unsupervised PCA analysis using phosphopeptides differentially expressed between MLLGA and MLLGB showed that new samples with KMT2A-MLLT4 and MLLT10 fusion proteins located close to MLLGA samples, whereas the sample with KMT2A-ELL/MLLT1 fusion located close to MLLGB samples (Figure 4a). Interestingly, KMTA2-MLLT3 and MLLT11 formed a new group with low PC2 that located much closer to MLLGA than to MLLGB (Figure 4a), suggesting that the phosphoproteomes of KMT2A-MLLT3 and MLLT11 samples are more similar to MLLGA than to MLLGB.

**Figure 4.**
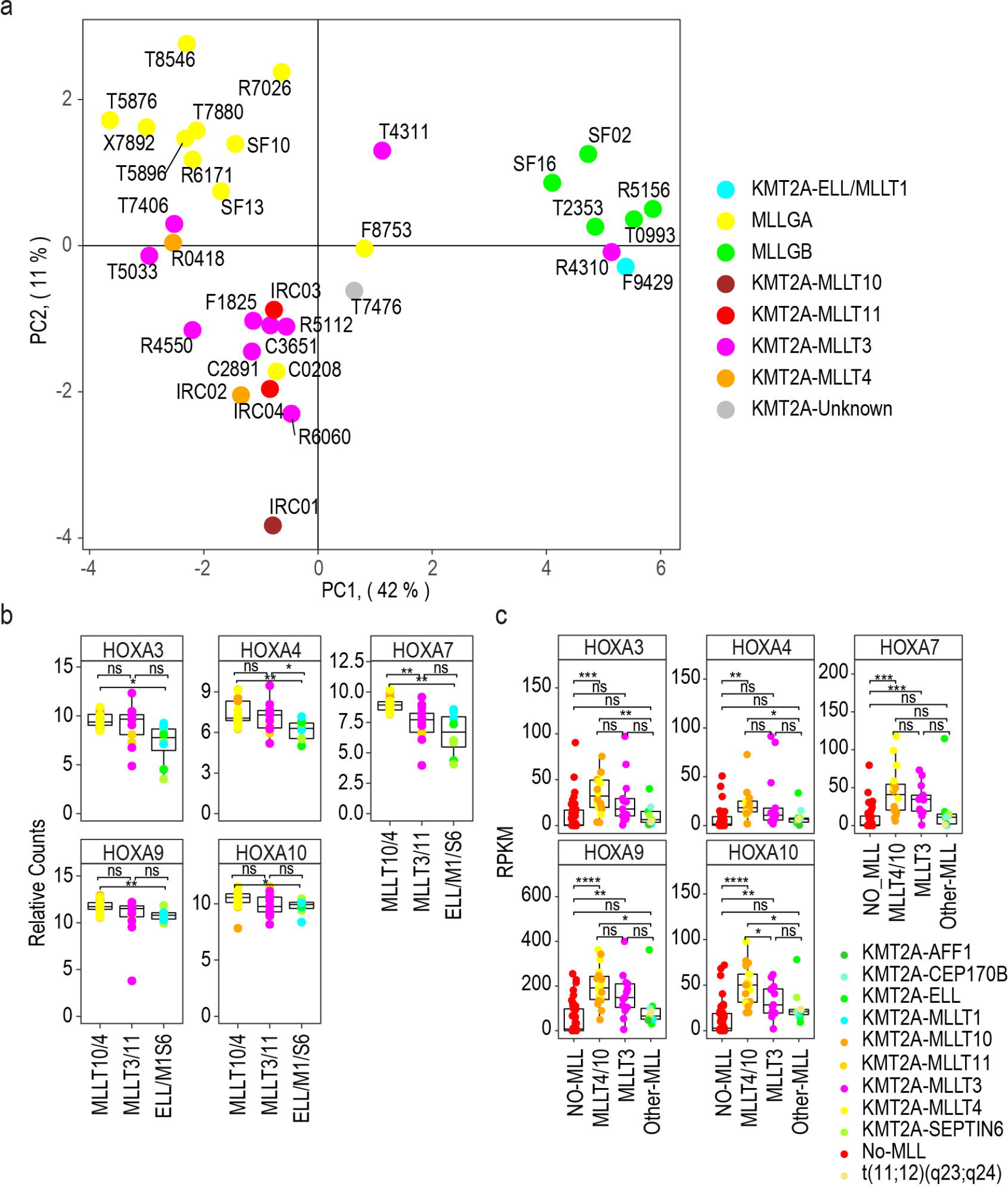
Phosphoproteome of KMT2A-MLL3 samples more similar to MLLGA than to MLLGB. **a** PCA patients with KMT2Ar-AML using differentially expressed phosphopeptides between MLLGA and MLLGB **b** HOXA gene expression in the dataset obtained from^30^. **c** HOXA gene expression in the dataset obtained from^31^. Data points represent individual patient observations. Boxplots indicate median, 1^st^ and 3^rd^ quartiles. Whiskers extends from the hinge to the largest and lowest value no further than 1.5 times the distance between the 1st and 3rd quartiles (**b** and **c**). For mRNA analysis, MLLT4/10/11 (n=23), MLLT3 (n=11), ELL/M1/S6 (n=8) in (**b**), and No-MLL (n=40), MLLT4/10 (n=18), MLLT3 (n=13), Other-MLL(n=9) in (**c**). Statistical significance was calculated using unpaired two-sided Student’s t-test. **** p ≤ 0.0001, *** p ≤ 0.001, ** p ≤ 0.01 and * p ≤ 0.05. (**b** and **c**).

To confirm our results showing differences in MLLGA and MLLGB in independent datasets, we collected publicly available mRNA expression data in a cohort of 42 young patients (aged <18 years) with KMT2Ar-AML from Pigazzi et al^30^. We considered cases with rearrangements involving *MLLT4* and *MLLT10* cytogenetically similar to our MLLGA and patients with rearrangements involving *ELL, MLLT1* and *SEPTIN6* similar to our MLLGB. Consistent with our results, we found that patients in the Pigazzi et al dataset with *MLLT4/MLLT10* rearrangements significantly overexpressed several HOXA genes when compared to patients with *ELL/MLLT1/SEPTIN6* (ELL/M1/S6) rearrangements (Figure 4b). MLLT3/MLLT11 patients showed higher expression of *HOXA4* than ELL/M1/S6 patients (Figure 4b). We also collected publicly available data for 80 patients of the TARGET-20 dataset^31^. We subdivided the dataset in patients with no *KMT2A* rearrangements (No-MLL) and patients with *KMT2A* rearrangements with *MLLT4* or *MLLT10* (MLLT4/10), *MLLT3* or other genes (Other-MLL). We found that MLLT4/10 patients overexpressed HOXA genes when compared to No-MLL and to Other-MLL patients (with the exception of *HOXA7*; Figure 4c). MLLT3 patients also overexpress *HOXA7*, *HOXA9* and *HOXA10* genes when compared to No-MLL, while no differences in HOXA gene expression was found between No-MLL and Other-MLL (Figure 4c). These data confirm our initial findings on the existence of two biochemically distinct KMT2Ar groups with distinct patterns of pathway activities and further suggest the presence of a third group comprising MLLT3 and MLLT11 that is closer to MLLGA than to MLLGB.

### MLLGA cases present an increased phosphorylation of proteins involved in RNA splicing, replication and DNA damage and an increased activity of CDK1

We next investigated whether, in addition to differences in HOXA gene expression and DOTL1/TEFb complexes, the MLLGA and MLLGB subgroups of KMT2Ar cases have other molecular differences. We found that MLLGA significantly increased or decreased the expression of multiple transcripts, proteins, phosphopeptides and acetylated or methylated peptides when compared to MLLGB or the No-MLL group (Figure 5a). AML groups also presented several hundred significant differences across protein or phosphorylated, methylated or acetylated peptide levels when compared to normal myeloid cells (Figure 5b).

**Figure 5.**
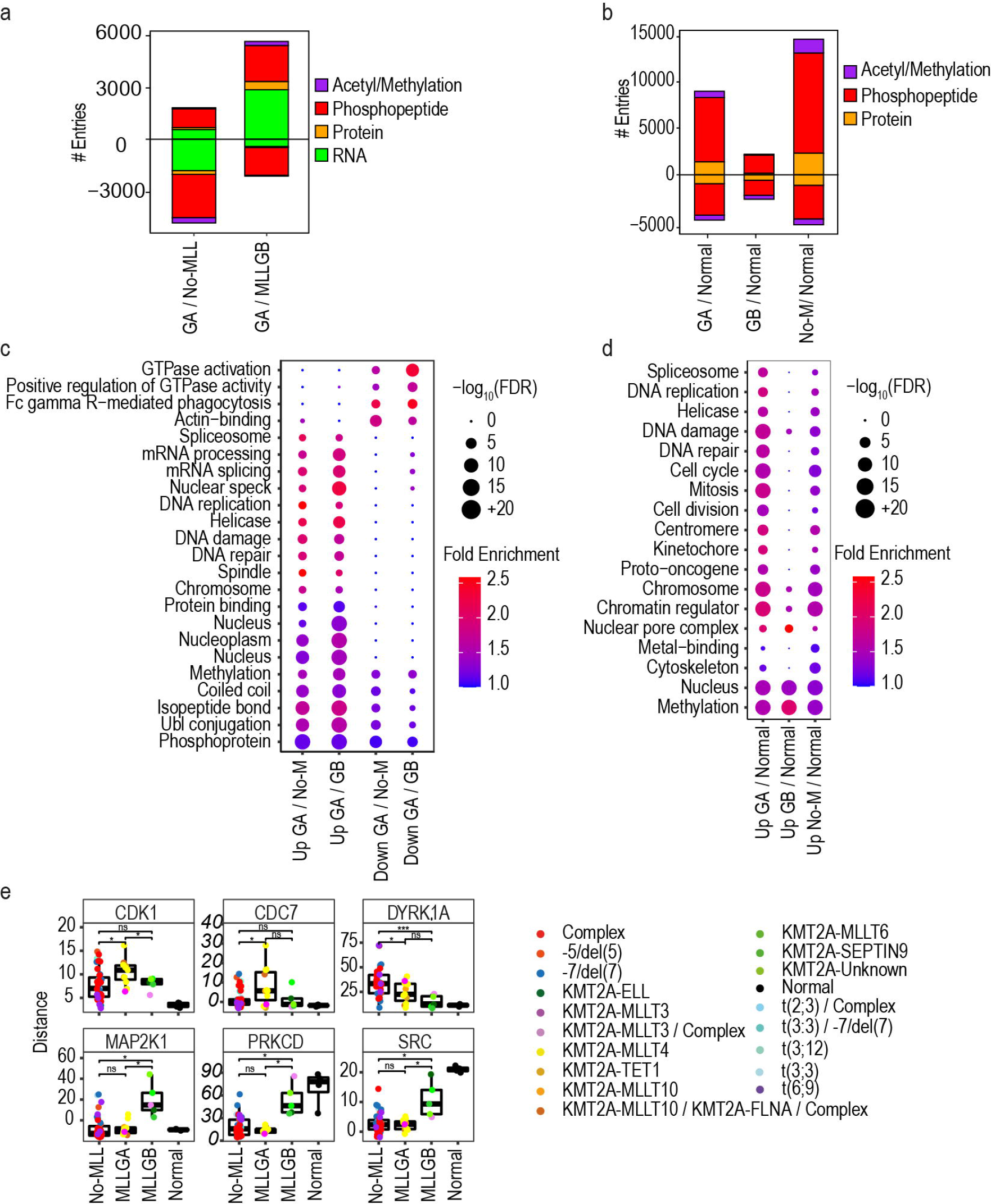
MLLGA increased the phosphorylation of proteins involved in RNA metabolism, replication and DNA damage response. **a** Expression levels of mRNA, protein, peptide phosphorylation and peptide methylation and acetylation across KMT2Ar groups. **b** Term enrichment analysis across KMT2Ar groups. **c** Estimation of kinase activity across KMT2Ar groups. **d** Expression levels of mRNA, protein, peptide phosphorylation and peptide methylation and acetylation in KMT2Ar groups compared to normal cells. **e** Term enrichment analysis in KMT2Ar groups compared to normal cells. Statistical significant was calculated using unpaired two-sided Student’s t-test. For mRNA analysis, Normal (n=0), MLLGA (n=9), MLLGB (n=3) and No-MLL (n=27) and for proteomics, phosphoproteomics, KSEA and methylation and acetylation analyses, Normal (n=4), MLLGA (n=11), MLLGB (n=5) and No-MLL (n=39) (**a** and **c-d**). Phosphopeptides were counted when p-value <0.05 and fold change (log2) > 0.7 (positive values) or <-0.7 (negative values) (**a** and **d**). Boxplots indicate median, 1^st^ and 3^rd^ quartiles. Whiskers extend from the hinge to the largest and lowest value no further than 1.5 times the distance between the 1st and 3rd quartiles. **** p ≤ 0.0001, *** p ≤ 0.001, ** p ≤ 0.01 and * p ≤ 0.05 (**c**). Statistical difference was calculated using a modified Fisher’s exact test. FDR values obtained by the adjustment of p-values using the Benjamini-Hochberg procedure (**b** and **e**). GA is MLLGA, GB is MLLGB and No-M is No-MLL.

Functional differences between groups of patients may be exploited to design therapies targeted to such subgroups. However, successful therapies should also aim to target functions that are not essential for the biology of normal cells. Ontology and pathway enrichment analysis on the phosphoproteomics dataset highlighted phosphoproteins linked to GTPase activity, actin binding and Fc receptor mediated phagocytosis to be significantly reduced in MLLGA relative to other cases (Figure 5c). Functions linked to RNA splicing, replication and the DNA damage response were enriched in the sets of phosphoproteins significantly increased in MLLGA compared to other cases (Figure 5c), and in MLLGA and No-MLL groups relative to myeloid cells from healthy donors (Normal) (Figure 5d), although the significance and the magnitude of the enrichment was higher in MLLGA cases (Figure 5d). DNA damage response proteins that significantly increased in phosphorylation in MLLGA included BRCA1 at S1466, S1542 and T1622, ERCC5 at S356, FANCE at S356, MDC1 at S168 and S196 and RIF1 at S22, among others (Supplementary Figure 5a). Similarly, the phosphorylation of proteins involved in DNA replication, such as RB1 at S37 and the MCM helicase components MCM2 at T106, MCM4 at S131 and MCM6 at T266, was also higher in MLLGA than in MLLGB, No-MLL and Normal groups (Supplementary Figure 5a and 5b). Together, these results uncovered biochemical pathways significantly enriched in MLLGA cells, including RNA splicing process, cell replication rate and DNA damage, when compared to the other AML subgroups and normal myeloid cells.

To identify differences in kinase activities across groups, we derived values of kinase activity from the phosphoproteomics data using KSEA^32^. The activities of kinases with roles in regulating mitosis and replication were increased in MLLGA. Specifically, MLLGA cases showed elevated CDK1 activity relative to all other groups, high CDC7 and CDK2 activities relative to No-MLL and Normal cases and high mTOR and other CDKs activities relative to Normal cases (Figure 5e and Supplementary Figure 5c). In addition, PRKCD and SRC activities were decreased in No-MLL and MLLGA when compared to MLLGB and Normal, while MAP2K1 activity was increased in MLLGB compared to all other groups (Figure 5e). Finally, several tyrosine kinases including SYK, EGFR, BTK, FYN and LCK were increased in the Normal group when compared to No-MLL and MLLGA (Supplementary Figure 5c). The No-MLL group presented and increased activity of DYRK1A when compared to the other two groups. These data show that MLLGA present a higher activity of kinases that positively regulate the cell cycle and cell proliferation when compared with normal myeloid cells and other AML groups.

The profound molecular differences between the KMT2Ar groups could translate into distinct clinical outcomes. A Kaplan-Maier curve showed that MLLGB cases trend to present a higher overall survival when compared to MLLGA and No-MLL although the difference was not statistically significant (Supplementary Figure 5d). Therefore, MLLGB cases trend to have a better prognosis than MLLGA and No-MLL patients.

### MLLGA is more sensitive to genotoxic drugs and inhibitors of mitotic kinases and IMPDH

We next aimed to assess whether differences in the biochemistry of KMT2Ar-AML subgroups translate into phenotypic differences that could be exploited clinically. Our multi-omics platform outlined above (Figure 1c) included a drug screening for 627 compounds based on the reduction of the cell number as a function of treatment, where high drug sensitivity scores (DSS) indicates high sensitivity to the compound. When considering the whole poor-risk and KMT2A-MLLT3 patient cohort, compounds that inhibit RNA synthesis and protein degradation were the most efficient agents on average, while receptor tyrosine kinase (RTK), PI3K/AKT/MTOR and epigenetic inhibitors where the most abundant compound types in our drug panel (Figure 6a).

**Figure 6.**
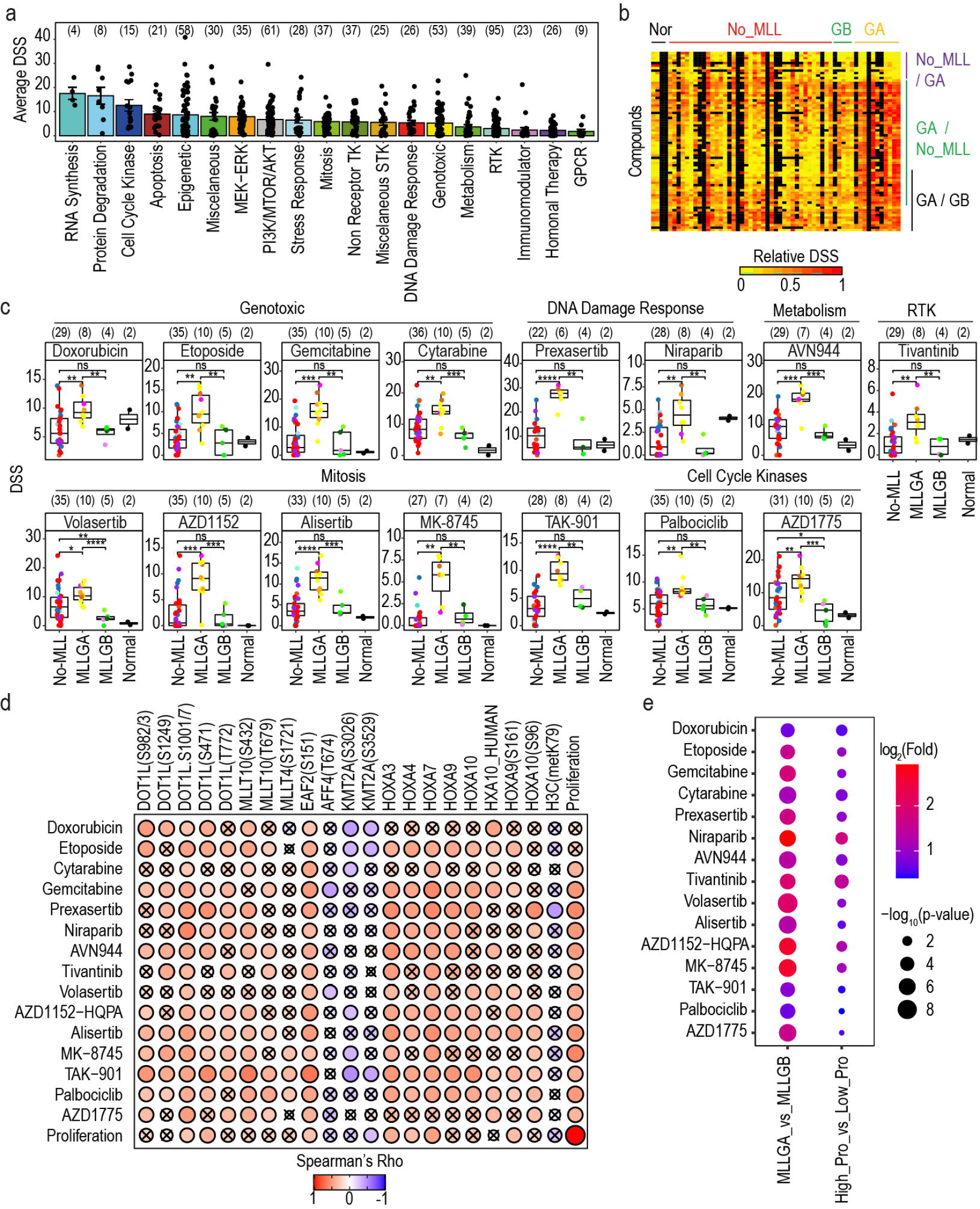
Genotoxic drugs and compounds targeting the DNA damage response and the cell cycle progression are highly effective in MLLGA. **a** Overview of the *ex vivo* response of AML samples to 482 compounds. **b** Compound response for agents that are more efficient in any of the KMT2Ar groups. **c** Compound response for agents that are more efficient in MLLGA than in MLLGB and No-MLL. **d** Spearman rank correlation rho values between compound response and proliferation or the mRNA, protein, phosphorylation or methylation/acetylation levels of DOT1L/TEFb complex components, HOXA genes or Histone H3. **e** Compound response (DSS) as a function of stratification based on KMT2Ar groups or proliferation for patients with KMT2Ar-AML. High pro group included samples with proliferation rate higher than the average rate for KMT2Ar-AML patients, while Low pro group included samples with proliferation rate lower than the average rate for KMT2Ar-AML patients. Only compounds tested in at least 30 patients were included, and data are represented as mean ± SEM (**a**). Dot colour indicate karyotype as in Figure 3b, and boxplots indicate median, 1^st^ and 3^rd^ quartiles. Whiskers extend from the hinge to the largest and lowest value no further than 1.5 times the distance between the 1st and 3rd quartiles (**c**). (n) is indicated in brackets (**a**, **c** and **e**). Statistical significance was calculated using unpaired two-sided Student’s t-test. **** p ≤ 0.0001, *** p ≤ 0.001, ** p ≤ 0.01 and * p ≤ 0.05. (**b** and **c**) Correlation was calculated using Spearman correlation and crosses denote correlations that are not statistically significant (p>0.5) (**d**).

Consistent with the increase in the activity of CDKs and pro-proliferative pathways in the MLLGA subgroup (Figure 5e), our drug screening showed mitotic poisons and genotoxic compounds particularly more effective in cells from MLLGA, relative to other cases. Specifically, 51 compounds were significantly more effective in MLLGA cases compared to those in the No-MLL group, while 17 agents, including apoptotic modulators like BCL-XL inhibitors, were significantly more effective in No-MLL (Figure 6b and Supplementary Figure 6a). Similarly, 25 compounds were significantly more effective in the MLLGA than in the MLLGB (Figure 6b and Supplementary Figure 6b). A total of 15 agents were significantly more effective in MLLGA when compared to both MLLGB and No-MLL (Figure 6b and c). These included topoisomerase inhibitors (etoposide and doxorubicin), nucleotide analogues (cytarabine and gemcitabine) and inhibitors developed to target CHEK1 (prexasertib), PARP1 (nilaparib), IMPDH (AVN-944), c-MET (tivatinib), PLK (volasertib), AURKA/B (AZD1152, alisertib, MK-8745 and TAK-901), CDK4/6 (palbociclib) and Wee1 (AZD1775) (Figure 6c). Correlation analysis revealed that the phosphorylation of several DOT1L/TEFb complex components and the expression of multiple HOXA genes were significantly associated with the responses to drugs that were more effective in MLLGA relative to other groups (Figure 6d). Interestingly, the phosphopeptides DOT1L(S1001/7) and EAF2(S159) presented a significantly positive correlation with the response to all these compounds (Figure 6d).

Since most of the 15 compounds that are more effective in MLLGA target mainly proliferative cells^33,34^, we then asked whether the rate of *ex vivo* cell proliferation determined responses to these agents. In agreement with these cases having high activity of the cell cycle related kinases CDK1 and CDC7 (Figure 5c), we found that cells from MLLGA samples proliferated significantly faster than those from other AML subgroups (Supplementary Figure 6c). Proliferation rate also positively correlated with the extent of phosphorylations of DOT1L and EAF2, the expression of HOXA3, 4 and 7 and the phosphorylation of HOXA9 (Figure 6d). Of note, cell proliferation rate significantly correlated with the DSS for 70 compounds (Supplementary Figure 6d) that included 13 of the 15 compounds that were more effective in MLLGA than in MLLGB and No-MLL (Figure 6d and Supplementary Figure 6e). However, when we compared classifications of KMT2Ar based on our phosphoproteomics signature or on the cellular proliferation rate, the MLLGA vs MLLGB stratification provided greater and more significant DSS differences than stratification by proliferation rate for all the MLLGA-specific compounds (Figure 6e). Together, these results uncovered 15 drugs that are particularly effective in MLLGA cases and indicate that, although contributing, proliferation rate is not the main determinant in how cells respond to MLLGA-specific compounds *ex vivo*.

### High expression of IMPDH2 and nucleolar proteins as determinants of IMPDH inhibitor efficacy in MLLGA

Having identified a set of drugs that are specific for the MLLGA subgroup of KMT2Ar-AML, we next explored factors, other than proliferation rate, that could be used to rationalise responses to these agents in the context of KMT2Ar-AML biology. We focused on AVN-944 because this compound inhibits the IMPDH enzymes that are essential for “de novo” synthesis of purine nucleotides^35^, and because we observed that AVN-944 DSS significantly correlates with IMPDH2 protein levels (Figure 7a, left panel), thus highlighting a clear link between drug activity and target expression. This correlation was similar to the correlation between the DSS for AVN-944 and the cellular proliferation rate (Supplementary Figure 6a, left panel). In a karyotype-based stratified analysis, IMPDH2 protein and RNA expression strongly correlated with AVN-944 DSS in KMT2Ar-AML samples, while no correlation was observed for samples with other poor-risk karyotypes (Figure 7a, right panel and b). In addition, AVN-994 DSS correlated better with IMPDH2 expression than with cell proliferation in KMT2Ar-AML samples (Supplementary Figure 7a, right panel). AVN-944 DSS did not correlate with IMPDH1 protein levels in KMT2Ar-AML, but it presented a strong negative correlation with the protein levels of DPYD, the limiting enzyme for the degradation of pyrimidine nucleotides^36^ (Supplementary Figure 7b and c). No association was observed between AVN-944 DSS and proliferation or the protein levels of IMPDH1 or DPYD in the No-MLL group (Supplementary Figure7 a-c). MLLGA, when compared to MLLGB, showed a higher sensitivity to AVN-944 and IMPDH2 expression and lower DPYD levels but presented no differences in proliferation and IMPDH1 expression (Supplementary Figure 7d and Figure 7c). These data suggest that IMPDH2 expression is a major determinant of IMPDH inhibitor sensitivity in KMT2Ar-AML and it is more expressed in MLLGA cases. In addition, since IMPDH and DPYD are key regulators of nucleotide metabolism, these data suggest that nucleotide metabolism is altered in MLLGA.

**Figure 7.**
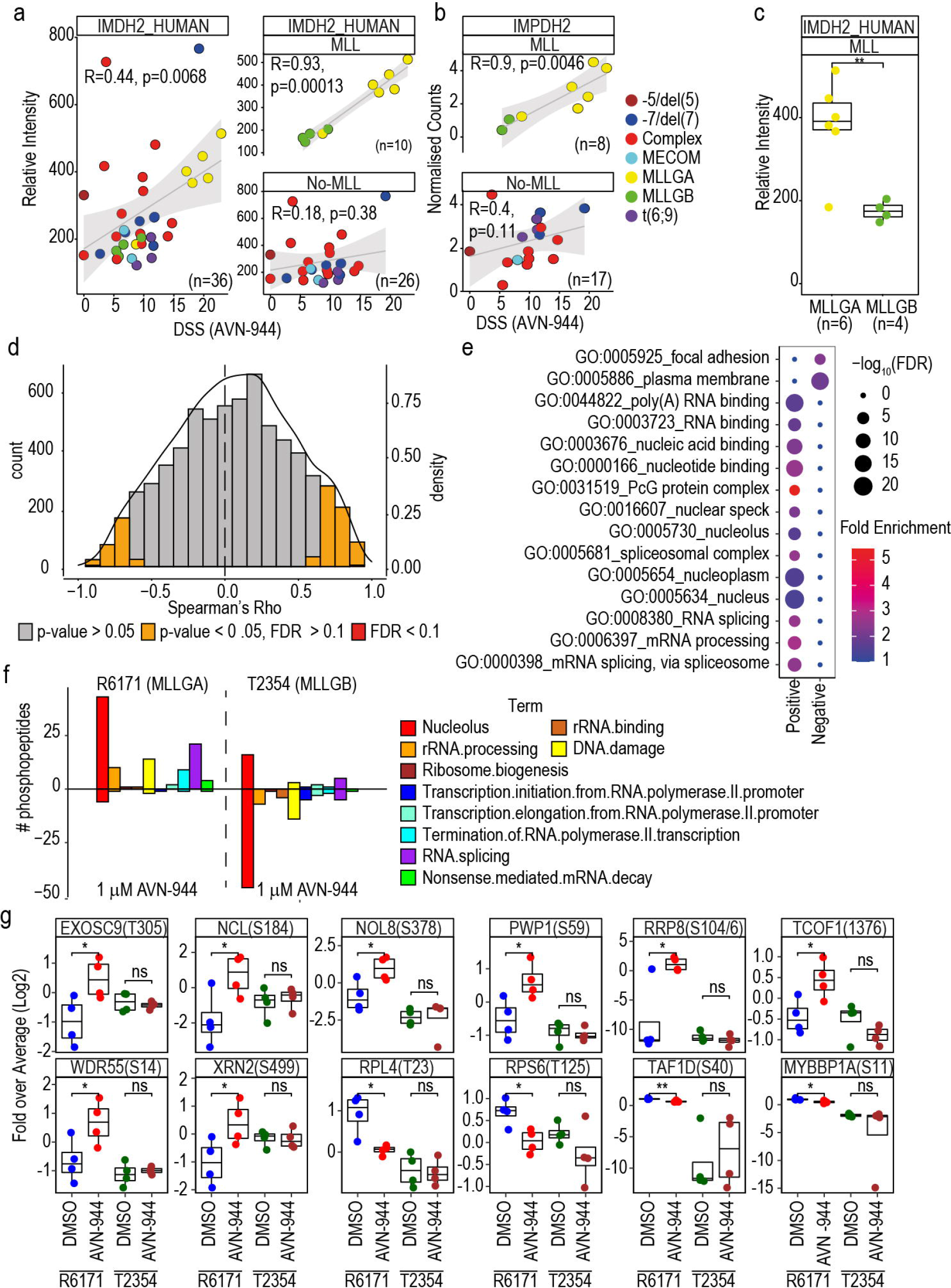
IMPDH2 expression and the nucleolar metabolism are significantly associated with responses to AVN-944 in KMT2Ar-AML. **a, b** Spearman rank correlation between response to AVN-944 and IMDH2 protein (**a**) and mRNA (**b**) expression. **c** Protein expression of IMPDH2 across MLLGA and MLLGB. **d** Distribution of Spearman rank correlation values between response (DSS) for AVN-944 and protein expression. **e** Term enrichment analysis in the sets of proteins significantly positively or negatively correlated in **d**. **f** Number of phosphopeptides significantly affected by 24h treatment with 1 μM AVN-944 as a function of term linkage. Phosphopeptides were counted when p-value <0.05 and fold change (log2) > 0.7 (positive values) or <-0.7 (negative values). **g** Selection of proteins significantly affected by treatment with 1 μM AVN-944 and linked to the nucleolus or the ribosome biology. Statistical significance was calculated using unpaired two-sided Student’s t-test (**c**, **f** and **g**). **** p ≤ 0.0001, *** p ≤ 0.001, ** p ≤ 0.01 and * p ≤ 0.05 (**c** and **g**). FDR values obtained by the adjustment of p-values using the Benjamini-Hochberg procedure (**c** and **d**). Statistical difference was calculated using a modified Fisher’s exact test (**d**). In **g**, n=4 biological replicates.

In addition to IMPDH2 and DPYD, sensitivity to AVN-944 positively or negatively correlated with the levels of 609 and 294 proteins, respectively (Figure 7d). Ontology and term enrichment analysis showed a significant enrichment of proteins linked to focal adhesion and the plasma membrane in the group that negatively correlated with the response to AVN-944 (Figure 7e). Conversely, proteins associated to RNA processing and the nucleolus were significantly enriched in the set that positively correlated with AVN-944 DSS (Figure 7e). Nucleolar proteins, whose expression correlated with AVN-944 sensitivity and were increased in MLLGA, included NPM1, NCL, UBF1 among others (Supplementary Figure 7e and f). These results show that proteins involved in nucleolar biology, which are highly expressed in MLLGA, are associated to the sensitivity to IMPDH inhibitors in KMT2Ar-AML.

Next, to stablish a causative link between high expressions of nucleolar proteins in MLLGA and high responses to IMPDH inhibitors seen in this KMT2Ar-AML subgroup, we investigated whether targeting IMPDH had a higher impact in the nucleolar activity in MLLGA than in MLLGB. We hypothesised that this greater interference would manifest as a differential phosphorylation of nucleolar proteins between MLLGA and MLLB treated with IMPDH inhibitors^37–39^. To this end, we analysed the phosphoproteome of samples R6171 and T2354, which were respectively classified as MLLGA and MLLGB, treated with AVN-944 for 24h. PCA of these data showed that R6171 and T2354 separated in PC1 space, while DMSO treated samples trended to separate from AVN-944 treated samples in the PC2 axis for R6171 (Supplementary Figure 8a). We also observed that AVN-944 significantly increased the phosphorylation of a similar number of peptides in R6171 and T2354 cells (Supplementary Figure 8b). More specifically, 1 μM AVN-994 significantly increased the phosphorylation of 43 phosphopeptides in proteins linked to the nucleolus in R6171, while it only increased the phosphorylation of 16 in T2354 (Figure 7f). Conversely, AVN-994 significantly reduced the phosphorylation of more peptides in T2354 than in R6171 (Supplementary Figure 8b). In the specific case of peptides in proteins linked to the nucleolus, AVN-994 reduced the phosphorylation of 46 peptides in T2354 and only 6 in R6171 (Figure 7f). Phosphopeptides located in proteins involved in the ribosome biology that were only significantly affected by AVN-944 in R6171 included EXOSC9, NCL, NOL8, PWP1, RPL4 and TAF1D among others (Figure 7g). After treatment with 1 μM AVN-944, the number of significantly increased phosphopeptides in proteins linked to RNA splicing and DNA damage was also higher in R6171 than in T2354 (Figure 7f). Overall, we found that the expression of key nucleolar regulators of rRNA synthesis and metabolism correlates with IMPDH inhibitor sensitivity. In addition, our results indicate that IMPDH inhibitors are, at least partially, more effective in MLLGA than in MLLGB because they have a greater impact on nucleolar biology in this KMT2Ar-AML subgroup.

## DISCUSSION

Recent advances in our understanding of AML biology and genetics has led to the development of new-targeted therapies to treat subpopulations of patients^40,41^. However, patients with poor-risk karyotypes still present low overall survival, thus highlighting the need for new drug targets and biomarkers to improve precision medicine in these patients^42^. Comprehensive analyses of the molecular pathways deregulated in malignant cells have provided substantial new knowledge of disease biology, from which new drug targets and signatures for patient stratification can be derived^16,19,21,43^. To understand the molecular and biochemical landscape of poor-risk AML, we performed an integrative analysis of data obtained from multiple omics platforms. Unlike previous multi-omics studies, we also carried out a comprehensive functional phenotypic analysis by drug sensitivity screening against 627 compounds (Figure 1 a-c, Supplementary Figure 1 and Supplementary Table 1)^44^. Detailed analysis of the other subgroups and data integration will be described elsewhere.

Here, we focused on the phosphoproteomics data of KMT2Ar cases because, interestingly, we found large differences in the phosphoproteomes of the different karyotypes within the poor-risk cases in our study, with KMT2Ar AML being the most biologically distinct AML subtype relative to other poor-risk cases (Figure 1e). Following this observation, we derived a phosphoproteomics signature that classified KMT2Ar-AML patients into two biochemically distinct groups, which we termed MLL group A (MLLGA) and MLL group B (MLLGB) (Figure 2 c and d). Of note, these results indicate that KMT2Ar-AML is a heterogeneous condition from the biochemical/proteomic standpoint.

Multiple KMT2A fusion proteins recruit DOT1L and TEFb complexes to the *HOXA* cluster and other KMT2A targets to promote transcription elongation by methylating histone H3 at K79 and phosphorylating the RNA polymerase II^12,13^. Aberrant expression of HOXA genes and other KMT2A targets promotes the leukaemogenic process^10^. In our study, MLLGA cases showed increased phosphorylation of several components of the DOT1L and TEFb complexes and elevated expression of multiple genes coded by the *HOXA* cluster when compared to other groups (Figure 3 a-c). HOXA gene expression has been used before to split leukaemia cases with KMT2A rearrangements. Indeed, ALL cases with KMT2A-AFF1 have been subdivided in two distinct subgroups based on HOXA gene expression with the group with low HOXA expression showing the worst prognosis^45–48^. However, our data suggest that the MLLGA that presented high HOXA gene expression show the worst overall survival in AML (Supplementary Figure 5d). In addition, MLLGA, but not MLLGB, cases presented a reduction of Histone H3 K79 methylation when compared to the No-MLL group (Figure 3d). Global reduction of histone H3 K79 methylation has been previously observed in KMT2Ar-AML patients^49^. Cases with KMT2A-MLLT3 rearrangements show much better outcome than the KMT2A-MLLT4 and MLLT10 cases represented in MLLGA and are not considered poor risk. However, these rearrangements generate fusion proteins able to recruit the DOT1L and TEFb complexes to the KMT2A target genes^29^. Our analysis show that KMT2A-MLLT3 cases together with KMT2A-MLLT11 cases are mainly represented in a third intermediate group that is more similar to MLLGA than to MLLGB (Figure 4a). To validate these results in two independent datasets based on RNA expression^30^, we used the KMT2A fusion partner to group KMT2Ar-AML cases into MLLGA, MLLGB and the intermediate MLLT3/MLLT11 group. Consistent with our data, we found that the groups similar to MLLGA over expressed multiple HOXA genes when compared to the groups similar to MLLGB while the MLLT3/MLLT11 group trend to express intermediate levels of these genes (Figure 4b and c). In summary, these data suggest that cells derived from MLLGA cases present a higher activity of the DOT1L and TEFb complexes recruited to the KMT2A target genes when compared to MLLGB and No-MLL, while cells from KMT2A-MLLT3 cases would present an intermediate activity more similar to MLLGA than to MLLGB.

Cells from MLLGA cases also showed an increased phosphorylation of proteins involved in RNA splicing, replication and DNA damage response (DDR) when compared to MLLGB, No-MLL and healthy myeloid cells (Figure 5c and d). CDC7 activity was increased only in MLLGA cases when compared to No-MLL (Figure 5e). CDC7 directly phosphorylates and regulates the activity of MCM, a helicase that plays a key role in replication^50^, and, consistently, we found that MLLGA presented an increased phosphorylation of three MCM subunits (Supplementary Figure 5b). Phosphorylation of key regulators of the DDR like the BRD1-BRCA1 complex or MDC1^51^ were also increased in MLLGA (Supplementary Figure 5a), which suggests that MLLGA cases are subjected to higher levels of DNA damage. It is tempting to speculate that the finding of increased phosphorylation of proteins that participate in the replication process may indicate that replicative stress could lead to an increase in DNA damage in these cases.

Since our phosphoproteomic signature stratified KMT2Ar-AML patients into two defined groups (Figures 2–5 and Supplementary Figures 4-5), we hypothesised that these subgroups would also present functional differences that could be exploited therapeutically. We therefore mined our *ex vivo* high content drug sensitivity screening against 627 compounds (Figure 6a)^52^. This analysis revealed that MLLGA samples, when compared to other groups, were more sensitive to 15 agents targeting topoisomerase II, nucleotide polymerization, CHEK1, PARP1, IMPDH2, c-Met, CDK4/6, Wee1, PLK and AURK (Figure 6b and c). Since most of these compounds target mainly proliferating cells^33,34^, we were prompted to investigate whether responses to these agents were correlated with the proliferation rate of the treated samples (Figure 6d and Supplementary Figure 5d-e). These experiments revealed that KMT2Ar-AML cases of the MLLGA, but not MLLGB, group proliferate at higher rate than those from the No-MLL group. Consistent with these observations, KSEA analysis of the phosphoproteomics data showed high activity of the mitotic kinase CDK1 in MLLGA cells (Figure 5e and Supplementary Figure 5c). However, our phosphoproteomics signature was more accurate than the proliferation rate in stratifying KMT2Ar-AML samples into responder and non-responder groups to these compounds (Figure 6e). Thus, these data indicate that the capacity of cells to proliferate influenced the response to the compounds that were more effective in MLLGA but other factors also contribute to the higher efficacy of these agents in this subgroup of KMT2Ar-AML cases.

MLLGA cases, represented by KMT2Ar involving MLLT4 and MLLT10 are highly sensitive to “ex vivo” treatment with the standard chemotherapeutics cytarabine and doxorubicin (Figure 5c). However, patients with these karyotypes present poorer outcomes when compared to other KMT2Ar (Supplementary Figure 5d)^53,54^. Discrepancies between outcome and “ex vivo” response to chemotherapy were also observed in infant ALL cases, where the presence KMT2Ar is associated with poor prognosis, but ALL cells with KMT2A-AFF1 rearrangements (the most frequent KMT2Ar in ALL) are more sensitive to “ex vitro” treatment with cytarabine^55,56^. In AML, high BM blast percentage is linked to KMT2Ar and low overall survival, but it is also a marker of increased proliferation, a major factor in the ex vivo response to chemotherapy^57,58^. Therefore, the results of our “ex vivo” assay for standard chemotherapy are in line with previous knowledge. As for the predictive nature of our “ex vivo” analysis, multiple publications have shown that the ex vivo drug response platform used in this study predicts patient response for several targeted drugs^52,59,60^. For example, a study of 37 relapsed AML patients showed a 59% objective response to the tailored therapies^61^. However, we acknowledge that it is difficult to extrapolate ex vivo response data to patient outcome in all cases. Our data indicate that MLLGA are more sensitive to chemotherapeutic drugs than an AML group that includes cases associated to high chemotherapy resistance (complex karyotype, TP53 mutations and t(6;9) rearrangements). In addition, KMT2A-MLLT3 cases who tend to respond to chemotherapy, are closer to MLLGA than MLLGB (Figure 4a) and would be expected to respond better to chemotherapeutics than No-MLL and MLLGB. These observations are, to some extent at least, compatible with the results of our ex vivo analysis, but clinical trials will ultimately be required to confirm the clinical relevance of our findings.

We focused on inhibitors of the inositol monophosphate dehydrogenase (IMPDH) because we found a clear link between target expression and drug efficacy within KMT2Ar-AML cases, and these compounds have been assessed for the treatment of cancer and other pathologies in multiple clinical trials; the FDA has consequently already approved their use in transplanted patients^35^. Consistent with our findings, a recent study reported that an IMPDH inhibitor reduced more effectively the proliferation in cord blood (CB) cells transformed with KMT2A-MLLT3 than in CB cells transformed with RUNX1-RUNX1T1 or in non-transformed CB cells^62^. IMPDHs are the rate limiting enzymes for *de novo* synthesis of guanosine, a precursor of the GMP required for DNA and RNA synthesis. Of the two IMPDH isoforms, only IMPDH2 has been found to be overexpressed in multiple cancer types^35^. We found that IMPDH2 but not IMPDH1 protein and RNA expression positively correlated with the response to the IMPDH inhibitor AVN-944 in KMT2Ar-AML cases but, of note, not in the No-MLL group (Figure 7 a-b and Supplementary Figure 7b). We also found that the expression of DPYD, the limiting enzyme for pyrimidine nucleotide degradation^36^, negatively correlated with the response to AVN-944 in KMT2Ar-AML (Supplementary Figure 7c). MLLGA responded better to AVN-944 than MLLGB, overexpressed IMPDH2 and presented lower levels of DPYD (Figure, 7c and Supplementary Figure 7d). Taken together, these data indicate that nucleotide metabolism is a determinant of responses to IMPDH inhibitors in KMT2Ar-AML but not in other AML cases.

In agreement with these results, we found that proteins whose expression positively correlated with AVN-944 response were enriched in nucleolar components. Anti-cancer properties of IMPDH inhibitors have been attributed to the inhibition of DNA synthesis^63^. However, recent work suggests that the increased guanosine synthesis in IMPDH2 overexpressing cells enables the formation of a pathologic nucleolus that uses the excess of guanosine for tRNA and rRNA synthesis and contributes to the malignant process. Of relevance, RNA processes and in particular rRNA synthesis utilize large amounts of nucleotides and generate vulnerabilities to IMPDH inhibitors. Therefore, cells that present a pathologic nucleolus are highly sensitive to IMPDH inhibitors^64–66^. Consistently, cells from MLLGA cases expressed significantly higher levels of key nucleolar proteins like nucleophosmin (NPM1), nucleolin (NCL) and UBF1 than MLLGB cases (Figure 7e and Supplementary Figure 7e-f). Furthermore, treatment with an IMPDH inhibitor impacted the phosphorylation of nucleolar proteins involved in rRNA expression and metabolism, like TAF1D and XRN2, and in the activation of TP53 after nucleolar stress, such as RPL4 and the MYBBP1-RPP8/NML axis in MLLGA but not in MLLGB (Figure 7f-g) ^67–70^. These data suggest that IMPDH inhibitors produced a greater impairment of ribosome biogenesis in MLLGA than in MLLGB. Therefore, our study indicates that the ability of IMPDH inhibitors to interfere with the nucleolar biology is, at least partially, responsible for the higher efficiency of these compounds in MLLGA.

In summary, we performed a deep multilayer molecular profiling of poor-risk and KMT2A-MLLT3 AML patients matched to the responses to hundreds of compounds in ex-vivo testing. Mining this dataset will allow identification of drug targets, mechanisms of drug action and biomarkers in poor-risk AML. As a proof of concept, here, we used these datasets to identify a phosphoproteomics signature that stratified KMT2Ar-AML cases in two biochemically and functionally distinct subgroups of patients. Although, ultimately, clinical trials will be required to confirm the clinical relevance of our findings, our study provides a rationale for the potential testing of IMPDH inhibitors (and potentially other mitotic and genotoxic drugs) in KMT2Ar-AML patients positive for the MLLGA phosphoproteomic signature identified in this study.

## MATERIALS AND METHODS

### Ethics Approval

Patients gave informed consent for the storage and use of their blood cells for research purposes. Experiments were performed in accordance with the Local Research Ethics Committee, as previously described^16^.

### Primary Samples

Mononuclear cells from peripheral blood or bone marrow biopsies were isolated in the BCI or FIMM tissue bank facilities and stored in liquid nitrogen.

### Proteomics, phosphoproteomics and PTM analysis

#### Thawing of AML primary samples

Cells were thawed at 37°C, transferred to 50 mL falcon tubes and incubated for 5 min at 37°C with 500 μL of DNAse Solution (Sigma Aldrich, Cat# D4513-1VL; resuspended in 10 mL of PBS). Then, 10 mL of 2% FBS in PBS were added and the cell suspension was centrifuged at 525g for 5 min at RT. Supernatant was discarded and cells were resuspended in 10 mL of complete IMDM (IMDM supplemented with 10% FBS and 1% Penicillin/Streptomycin; Thermo-Fisher Scientific Cat# 12440053, 10500-064 and 15140122, respectively), filtered through a 70 μm strainer and counted. Dilutions of 15×10^6^ cells in 10 mL of complete IMDM for each sample were incubated for 3h in an incubator at 37°C and 5% CO_2_. For cell harvesting, cell suspensions were centrifuged for 5 min at 525g at 5°C, pellets were washed twice with PBS supplemented with phosphatase inhibitors (1 mM Na_3_VO_4_ and 1 mM NaF). Pellets were transferred to low protein binding tubes (Sigma-Aldrich, Cat# Z666513-100EA) and stored at −80°C

#### Purification of myeloid cells from PBMCs

Myeloid cells were purified from PBMCs using Easy Sep Human Myeloid positive selection kit (Stem cell technology; Cat# 18653). The kit positively selects myeloid cells positive for CD33 and CD66b. detailed explanation of procedure can be found in the supplementary materials.

#### ANV-944 treatment

R6171 and T3254 were thawed as described for thawing AML primary samples. 10 to 15×10^6^ cells were resuspended in 10 mL of complete IMDM and treated with either vehicle (DMSO) or the indicated concentrations of AVN-944 (MedChemExpress, Cat# HY-13560) for 24h. The final concentration of DMSO (Thermo-Fisher Scientific, Cat# 10213810) was maintained at 0.1%. Cells were harvested as indicated for thawing AML primary samples and processed for phosphoproteomics analysis using 250 μg of protein.

#### Sample processing and MS analysis

Samples were processed and analysed as previously described^16,32^. For the poor risk cohort, samples were lysed and 250 μg of protein were subjected to reduction, alkylation and trypsin digestion. Protein extracts from two healthy donors were pooled to obtain 3 samples with 250 μg of protein and the fourth sample was obtained by pooling protein extracts from all 6 donors. After digestion, 220 μg were used for phoshoproteomics analysis and 30 μg for proteomics analysis. Samples for phosphoproteomics analysis were subjected to desalting using Oasis cartridges and phosphoenrichment using TiO_2_, while samples for proteomics were desalted using carbon spin tips. Samples were dried in a speed vac, resuspended in reconstitution buffer and run in a LC-MS/MS platform. For the KMT2Ar cohort, 100μg of protein were subjected to reduction, alkylation and trypsin digestion. Peptide suspensions were subjected to desalting and phosphoenrichment using TiO_2_. The LC-MS/MS platform consisted of a Dionex UltiMate 3000 RSLC coupled to Q Exactive™ Plus Orbitrap Mass Spectrometer (Thermo-Fisher Scientific) through an EASY-Spray source. Peptides and proteins were identified using Mascot (v2.6.0) and quantified using Pescal (vBeta2018). A detailed explanation for the generation and analysis of mass spectrometry data can be found as supplementary material

### DNA Sequencing

Total RNA and genomic DNA were extracted from patient PBMC or BM samples using the RNeasy and DNeasy Blood & Tissue Kits (Qiagen), respectively, following standard procedures, and concentrations were determined using Qubit® 3.0 Fluorometer.

DNA targeted next-generation sequencing (NGS) analysis was performed using the TruSight Myeloid Sequencing panel (Illumina, San Diego, CA, USA) targeting 54 genes (full coding exons of 15 genes: BCOR, BCORL1, CDKN2A, CEBPA, CUX1, DNMT3A, ETV6/TEL, EZH2, KDM6A, IKZF1, PHF6, RAD21, RUNX1/AML1, STAG2, ZRSR2, and exonic hotspots of 39 genes: ABL1, ASXL1, ATRX, BRAF, CALR, CBL, CBLB, CBLC, CSF3R, FBXW7, FLT3, GATA1, GATA2, GNAS, HRAS, IDH1, IDH2, JAK2, JAK3, KIT, KRAS, KMT2A/MLL, MPL, MYD88, NOTCH1, NPM1, NRAS, PDGFRA, PTEN, PTPN11, SETBP1, SF3B1, SMC1A, SMC3, SRSF2, TET2, TP53, U2AF1, WT1). Amplicon sequencing libraries were prepared from 50 ng of DNA per sample. Input DNA quantitation was performed using a Qubit 3.0 Fluorometer with Qubit 1X dsDNA HS Assay Kit (Life Technologies). After quality control and equimolar pooling paired-end sequencing of the libraries was performed on a NextSeq (Illumina, San Diego, CA, USA) instrument with NextSeq 500 High Output v2 Kit to generate 2×150 read lengths according to manufacturer’s instructions. Sequence data were analyzed using the TruSeq Amplicon v3.0.0 app in BaseSpace™ Sequence Hub. After demultiplexing and FASTQ file generation, the software uses a custom banded Smith-Waterman aligner to align the reads against the human hg19 reference genome to create BAM files. Variant calling for the specified regions was performed using the Somatic Variant Caller (5% threshold, read stitching on).

### mRNA Sequencing

RNA libraries were prepared using DNBseq sequencing technology and sequenced on a BGISEQ-500 sequencer generating 2×100bp paired-end reads. RNA-seq was performed at a depth of 100 million reads per sample. RNA-Seq data was aligned using HiSat2 v2.1.0 to GRCh38.p10 with the ensembl v91 reference annotation. Gene level counts were generated using htseq-count v0.13.5

### Drug sensitivity and resistance testing (DSRT)

A library of 627 commercially available chemotherapeutic and targeted oncology compounds were tested at 5 concentrations in 10-fold dilutions. The library consisted of 180 approved drugs, 334 investigational compounds and 113 probes (Supplementary Table X). The chemical compounds, DMSO (negative control) and benzethonium chloride (positive controls) were added to 384-well plates using an acoustic liquid dispensing system ECHO 500/550 (Labcyte). Biobanked frozen MNCs were thawed, resuspended in CM (conditioned medium) constituted of 77.5% RPMI 1640, 10% FCS, 12.5% human HS-5 bone marrow stromal cell line derived conditioned medium and 1% penicillin and streptomycin, let recover for 3h, and live cells counted. Compounds were first dissolved by adding 5μL of cell free medium, followed by 20μL cell suspension containing 5,000 viable cells to each well using EL 406 plate washer-dispenser (BioTek). The plates were incubated at 37°C in 5% CO_2_ for 72h. Subsequently, CellTiter-Glo (Promega) reagent was added to all wells and cell viability was measured using a PHERA star FS multimode plate reader (BMG Labtech).

The drug responses passing the data quality assessment were included in further analysis^71^. Drug sensitivity scores (DSS) were calculated as shown previously^44^.

### Estimation of the Proliferation rate

Proliferation rate was estimated form drug sensitivity screening data using the change in luminescence of untreated cells between day 0 and day 3 of treatment. Thus, Luminescence ratio = (luminescence at day 0/luminescence at day3)×100.

### Statistics

Statistical analysis was carried out in excel or in R-4.0.0 using base functions or the ggpubr package (https://CRAN.R-project.org/package=ggpubr). ML model performance was evaluated using caret package and base functions (https://CRAN.R-project.org/package=caret). Correlation matrices were generated using the corrplot package (https://CRAN.R-project.org/package=corrplot) and cluster dendrograms were generated using factoextra package (https://cran.r-project.org/web/packages/factoextra/index.html). Term enrichment analysis of protein subsets was performed using David Bioinformatics (https://david.ncifcrf.gov/). Details of the procedure can be found as supplementary material. David Bioinformatics results were parsed to CSV files and dot plots were constructed in an R environment using the ggplot2 package (https://CRAN.R-project.org/package=ggplot2).

## DATA AVAILABILITY

Raw mass spectrometry, mRNA sequencing and the DNA sequencing data are available upon request.

## ACKNOWLEDGEMENTS

We thank Adrian Kontor for technical help with the manipulation of AML primary samples, Sarah Mueller for managing the supply of AML primary samples, Janet Matthews for assisting with the processing of patient clinical data, Ruth Osuntola for technical assistance with the mass spectrometry experiments and the FIMM High Throughput Biomedicine Unit for their expert technical support. This work was mainly funded by Cancer Research UK (C15966/A24375) with additional contribution from Blood Cancer UK (20008). All authors have read and approved the article.

## CONTRIBUTIONS

P.C. and P.R.C. designed experiments, analysed the data and wrote the paper. P.C., A.R.M., S.K., V.R., K.R.P., C.B., J.J.M., C.H., F.B.C. and A.P. performed experiments and analysed data. P.C. and P.R.C. integrated data. J.G., F.M.M., D.C.T. and C.H. designed experiments and provided patient samples. P.R.C. and J.F. conceptualized the study. All authors contributed to writing the manuscript.

