## Supplementary Figure for "Integrative phosphoproteomics defines two biologically distinct groups of KMT2A rearranged acute myeloid leukaemia with different drug response phenotypes"

**This PDF file includes:**

Materials and Methods

Figures. S1 to S8

Materials and Methods

Purification of myeloid cells from PBMCs

Myeloid cells were purified from PBMCs using Easy Sep Human Myeloid positive selection kit (Stem cell technology; Cat# 18653). The kit positively selects myeloid cells positive for CD33 and CD66b. Cells from 6 donors were thawed, counted and centrifuged 1500 rpms for 5 min at RT. Cell pellets were resuspended in 1 mL of antibody solution (10% antibody mix, 2% FBS, 1% Penicillin/Streptomycin and 1% DNAse solution in PBS) and incubated for 15 min. Then, 100 uL of bead solution were added, cell suspensions were incubated for 10 min and transferred to polystyrene FACS tubes. Tubes were inserted into magnets and incubated for 5 min. Cell solution was decanted, tubes were taken out of the magnets and cells were resuspended in 2.5 mL of FBS solution (2% FBS, 1% Penicillin/Streptomycin and 1% DNAse solution in PBS). Tubes were inserted in the magnet, incubated for 5 min and the solution was decanted. Cells were resuspended in 10 mL of complete IMDM, incubated for 3h at 37°C with 5% CO_2_ and processed for mass spectrometry analysis as described for AML samples. When protein was quantified samples from 6 donors were pooled to generate 4 healthy donor samples with 250 µg. Samples were pooled as follows: Sample 1: Donors A and B (50% each); Sample 2 Donors C and D (50% each); Sample 3: Donors E and F (50% each); Sample 4: Donors A to F (variable % each)

Sample preparation for phosphoproteomics and proteomics analysis

Phosphoproteomics and proteomics analysis was carried out as previously described. AML cell pellets were lysed in 300 µL of urea buffer (8 M urea in 20 mM HEPES, pH: 8.0, supplemented with 1 mM Na_3_VO_4_, 1 mM NaF, 1 mM Na_4_P_2_O_7_ and 1 mM β-glycerophosphate). AML cell lysates were homogenised by sonication for 90 cycles (30 s on 30 s off) in a Diagenode Bioruptor® Plus and insoluble material was removed by centrifugation. Proteins were quantified using BCA protein assay. Then, 250 μg of protein were subjected to cysteine reduction and alkylation using sequential incubation with 10 mM dithiothreitol (DDT) and 16.6 mM iodoacetamide (IAM) for 1 h and 30 min, respectively, at 25°C with agitation. Trypsin beads (50% slurry of TLCK-trypsin, Thermo-Fisher Scientific; Cat. #20230) were equilibrated with 3 washes with 20 mM HEPES (pH 8.0), the urea concentration in the protein suspensions was reduced to 2 M by the addition of 900 µL of 20 mM HEPES (pH 8.0), 100 μL of equilibrated trypsin beads were added and samples were incubated overnight at 37°C. Trypsin beads were removed by centrifugation (2000 xg at 5°C for 5 min) and samples were divided in 220 µg for phosphoproteomics analysis and 30 µg (200 µL) for proteomics analysis.

For phosphoproteomics analysis, peptide solutions were desalted using Oasis HLB cartridges (Waters) following the manufacturer’s indications. Briefly, cartridges were set in a vacuum manifold device and the pressure was adjusted to 5 mmHg. Then, cartridges were conditioned with 1 mL acetonitrile (ACN) and equilibrated with 1.5 mL of wash solution (0.1% trifluoroacetic acid (TFA), 2% ACN). Peptides were loaded in the cartridges and washed twice with 1 mL of wash solution. Finally, peptides were eluted with 500 µL of glycolic acid buffer 1 (1 M glycolic acid, 5% TFA, 50% ACN). Enrichment of phosphorylated peptides was performed with TiO_2_ Beads (GL Sciences). Desalting eluents were normalized to 1 mL with glycolic acid buffer 2 (1 M glycolic acid, 5% TFA, 80% ACN) and incubated with 50 µl of TiO_2_ buffer (500 mg TiO_2_ beads in 1 mL of 1% TFA) for 5 min at room temperature. TiO_2_ beads were packed by centrifugation into empty spin columns previously washed with ACN. TiO_2_ beads were sequentially washed by centrifugation (1500 xg for 3 min) with 100 µL of glycolic acid buffer 2, ammonium acetate solution (100 mM ammonium acetate in 25% ACN) and twice with neutral solution (10% ACN). For phosphopeptide elution, spin tips were transferred to fresh tubes, 50 µL of elution solution 1 (5% NH_4_OH, 7.5% ACN) were added and tips were centrifuged at 1500 xg for 3 min. The elution step was repeated a total of 4 times. Finally, samples were snap frozen in dry ice, dried in a SpeedVac vacuum concentrator and phosphopeptide pellets were stored at −80°C.

For proteomics experiments, peptide solutions were desalted using C18+carbon top tips (Glygen Corporation). The tips were conditioned twice with 200 µL of elution solution 2 (70/30 ACN/ H2O + 0.1% TFA) and equilibrated twice with 200 µL of wash solution. Samples were loaded into the top tips and washed twice with 200 µL of wash solution. For peptide elution, the tips were transferred to fresh tubes and peptides were eluted four times with 50 µL of elution solution 2. Tips were centrifuged at 1500 xg for 5 min at 5˚C in all desalting steps. Eluted peptide solutions were dried in a SpeedVac vacuum concentrator and peptide pellets were stored at -80 ˚C.

Mass spectrometry

Mass spectrometry for identification and quantification of proteins and phosphopeptides was carried out by LC-MS/MS as described before^1-3^. For phosphoproteomics analysis, peptide pellets were reconstituted in 8 µL of reconstitution buffer (20 fmol/µL enolase in 3% ACN, 0.1% TFA) and 5 µL were loaded onto an LC-MS/MS system. For proteomics analysis, peptide pellets were reconstituted in 8 µL of 0.1% TFA, 2 µL of this solution were further diluted in 8 µL of reconstitution buffer and 2 µL were injected into the LC-MS/MS system.

The LC-MS/MS platform consisted of a Dionex UltiMate 3000 RSLC coupled to Q Exactive™ Plus Orbitrap Mass Spectrometer (Thermo Fisher Scientific) through an EASY-Spray source. Mobile phases for the chromatographic separation of the peptides consisted of Solvent A (3% ACN; 0.1% FA) and Solvent B (99.9% ACN; 0.1% FA). Peptides were loaded in a μ-pre-column and separated in an analytical column using a gradient running from 3% to 23% B over 60 min (for phosphoproteomics) or 120 min (for proteomics). The UPLC system delivered a flow of 2 µL/min (loading) and 250 nL/min (gradient elution). The Q Exactive Plus operated a duty cycle of 2.1s. Thus, it acquired full scan survey spectra (m/z 375–1500) with a 70,000 FWHM resolution followed by data-dependent acquisition in which the 15 most intense ions were selected for HCD (higher energy collisional dissociation) and MS/MS scanning (200–2000 m/z) with a resolution of 17,500 FWHM. A dynamic exclusion period of 30s was enabled with m/z window of ±10 ppm.

Identification of proteins and phosphorylated, acetylated and methylated peptides

Peptide identification from MS data was automated using Mascot Daemon (v2.6.0) workflow in which Mascot Distiller (v2.6.1.0) generated peak list files (MGFs) from RAW data and the Mascot search engine (v2.6) matched the MS/MS data stored in the MGF files to peptides using the SwissProt Database restricted to the *Homo sapiens* taxon (SwissProt_Latest_2018_09.fasta; 20411 sequences). Searches had a FDR of ~1% and allowed 2 trypsin missed cleavages, mass tolerance of ±10 ppm for the MS scans and ±25 mmu for the MS/MS scans, carbamidomethyl Cys as a fixed modification and PyroGlu on N-terminal Gln, oxidation of Met and phosphorylation on Ser, Thr, and Tyr as variable modifications (phosphorylation was only included for searches performed using phosphoproteomics data). Identification of peptides with PTMs was performed by searching MGF files generated from proteomics RAW files using the parameters above mentioning and allowing methyl R, methyl K, dimethyl K, trimethyl K and acetyl K as variable modifications. Searches in R and K were performed independently.

Phosphopeptide and protein quantification

Pescal software quantified the identified peptides using a label free procedure based on extracted ion chromatograms (XICs). Missing data points were minimized by constructing XICs across all LC-MS/MS runs for all the peptides identified in at least one of the LC-MS/MS runs^4^. XIC mass and retention time windows were ±7 ppm and ±2 min, respectively. Quantification of peptides was achieved by measuring the area under the peak of the XICs.

Data source and processing

RNA-Seq data was generated in-house or obtained from^5^. Mass spectrometry, drug response and panel sequencing data was generated in-house. For mass spectrometry datasets, individual peptide intensity values in each sample were normalized to the sum of the intensity values of all the peptides quantified in that sample. Data points not quantified for a particular peptide were given a peptide intensity value equal to the minimum intensity value quantified in the sample divided by 10. For phosphoproteomics experiments, we obtained a phosphorylation index (ppIndex) by summing the signals of all peptide ions containing the same modification site. For the proteomics experiment, protein intensity values were calculated by adding the intensities of all the peptides derived from a protein. Protein score values were expressed as the maximum Mascot protein score value obtained across samples. Drug response data were subjected to a quality control and drug responses passing the data quality assessment were included in further analysis^6^. Drug sensitivity scores (DSS) were calculated as shown previously^7^

Term enrichment analysis

Term enrichment analysis of protein subsets was performed using David Bioinformatics (<https://david.ncifcrf.gov/>). Enrichment was calculated as *[a/b]/[c/d]*, were *a* is number of proteins in the subset of interest that belong to a given term, *b* is number of proteins in the subset of interest, *c* is number of proteins that belong to such term in the background data and *d* is the size of the background data. All proteins identified in the dataset of interest were used as background. The p-values were obtained by a modified Fisher’s exact test and adjusted using the FDR method. David Bioinformatics results were parsed to CSV files and dot plots were constructed in an R environment using the ggplot2 package ([https://CRAN.R-project.org/package=ggplot2](about:blank)).

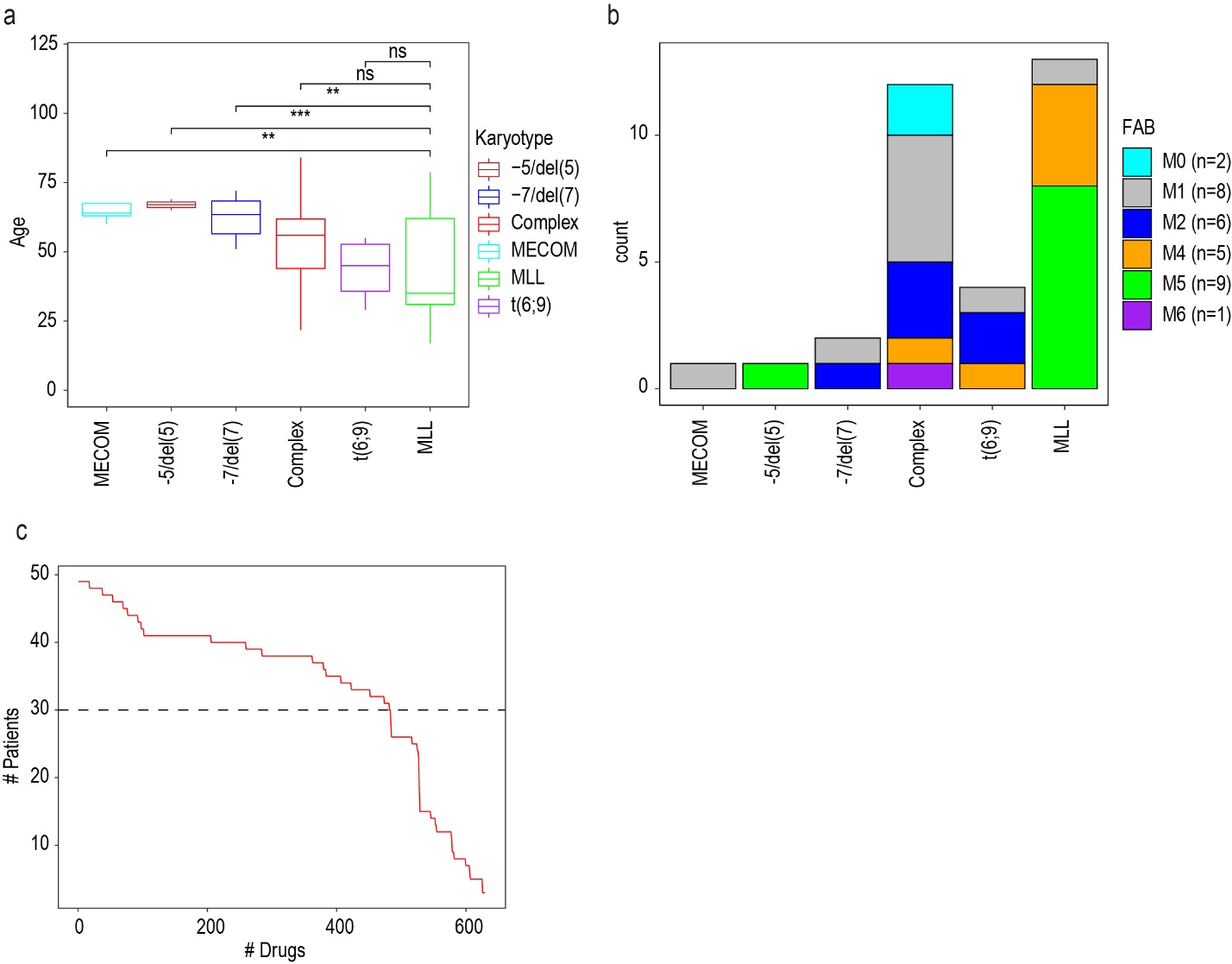

Figure. S1.

**Clinical features of the cohort of AML patients and number compounds used in the ex vivo drug screen**. **a** Age of AML patients as a function of karyotype. **b** FAB subtype classification of leukemic cells from AML patients as a function of karyotype. **c** Number of compounds tested in the ex vivo drug screen per patient. Boxplots indicate median, 1^st^ and 3^rd^ quartiles. Whiskers extends from the hinge to the largest and lowest value no further than 1.5 times the distance between the 1st and 3rd quartiles. Statistical significant was calculated using unpaired two-sided Student’s t-test. **** p ≤ 0.0001, *** p ≤ 0.001, ** p ≤ 0.01 and * p ≤ 0.05 (**b**).

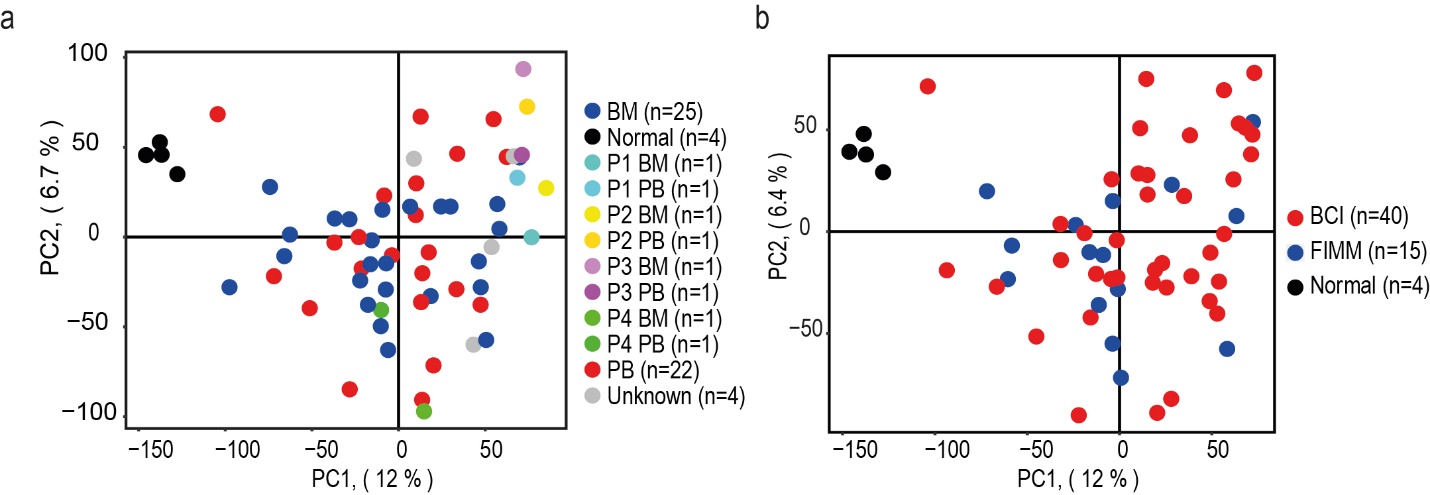

Figure. S2.

**Separation of samples as a function of source and origin using phosphoproteomics data.** **a** PCA using phosphoproteomics data to assess sample separation based on source. **b** PCA using phosphoproteomics data to assess sample separation based on origin. BM indicates bone Marron and PB indicates peripheral blood. Paired samples for 4 patients (P1-P4) are specifically colour coded.

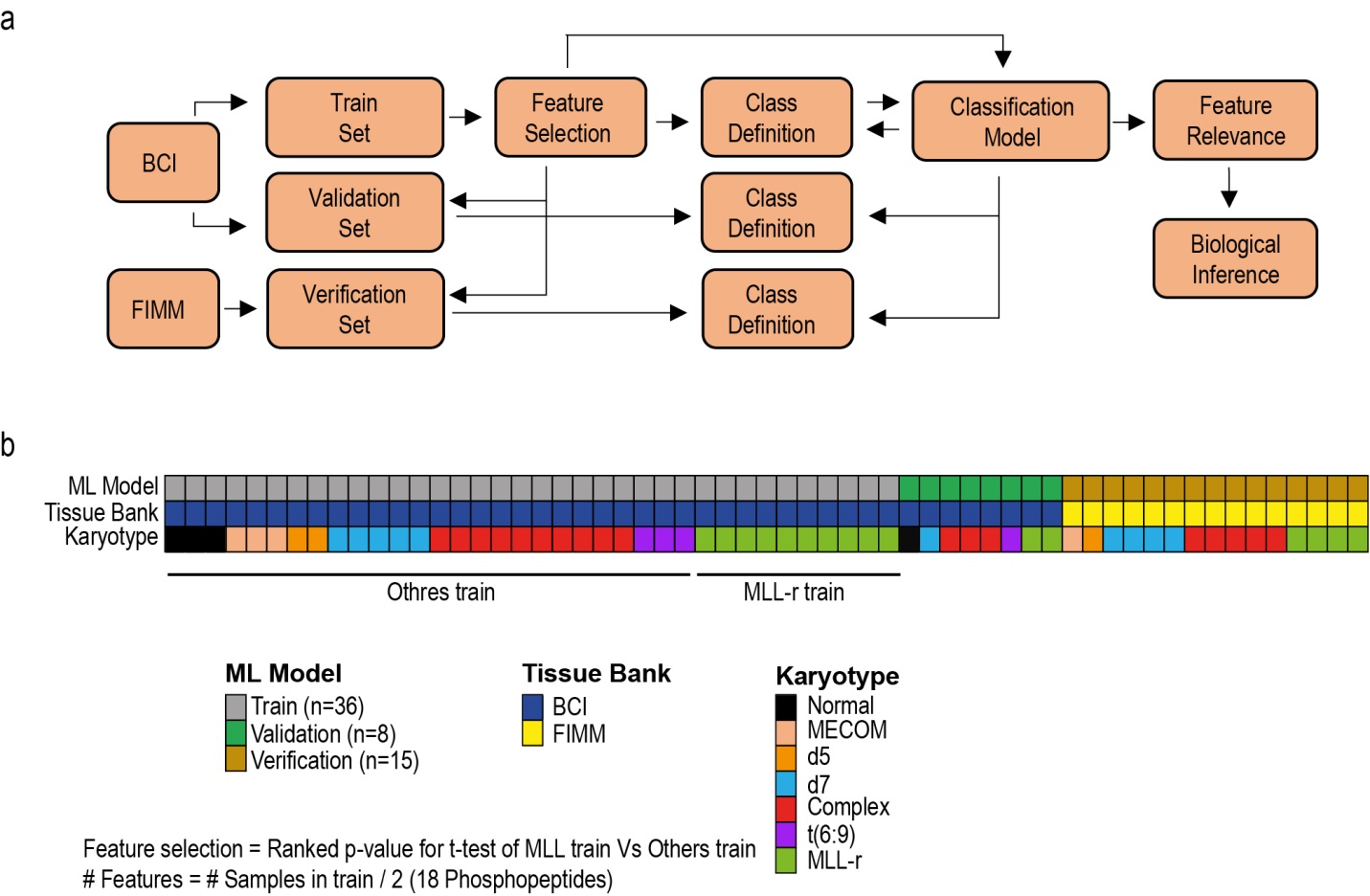

Figure. S3.

**Machine learning (ML) workflow followed to generate a phosphoprotemics signature and use it to stratify AML samples based on KMT2A rearrangements. ­a** Scheme of the machine learning workflow. b Samples compared in the t-test used to select the phosphopeptides comprised in the phosphoproteomics signature for KMT2Ar.

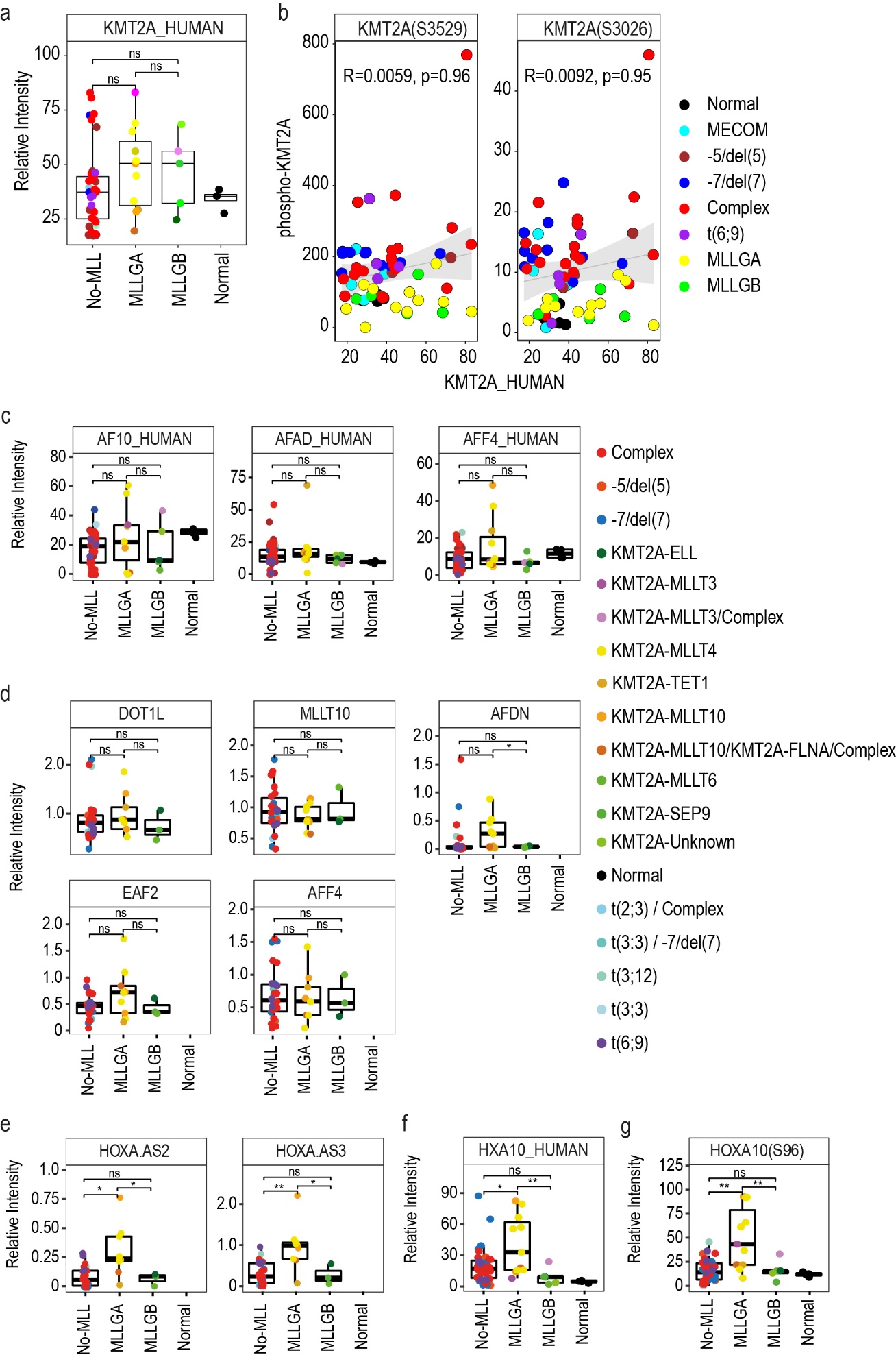

Figure. S4.

**Expression or phosphorylation of proteins or RNAs related to the biology of KMT2A fusion proteins as a function of KMT2A rearranged group.** **a** Protein expression of KMT2A. **b** Spearman rank correlation between protein expression and phosphorylation of KMT2A. **c** Protein expression of the genes MLLT10 (AF10), MLLT4 (AFAD) and AFF4 as a function of KMT2A rearranged group. **d** mRNA expression of DOT1L or TEFb complex components as a function of KMT2A rearranged group. **e** Expression of long non-coding RNAs coded in the HOXA cluster as a function of KMT2A rearranged group. **f** Protein expression of HOXA10 (HXA10) as a function of KMT2A rearranged group. **g** Phosphorylation of HOXA10 as a function of KMT2A rearranged group. Boxplots indicate median, 1^st^ and 3^rd^ quartiles. Whiskers extends from the hinge to the largest and lowest value no further than 1.5 times the distance between the 1st and 3rd quartiles. Statistical significance was calculated using unpaired two-sided Student’s t-test (**a** and **c**-**g**). **** p ≤ 0.0001, *** p ≤ 0.001, ** p ≤ 0.01 and * p ≤ 0.05. For mRNA analysis (**e**), Normal (n=0), MLLGA (n=9), MLLGB (n=3) and No-MLL (n=27) and for protein and phosphoproteomics analyses (**a, c**, **d,** **f** and **g**), Normal (n=4), MLLGA (n=11), MLLGB (n=5) and No-MLL (n=39); (n=59) for (**b**).

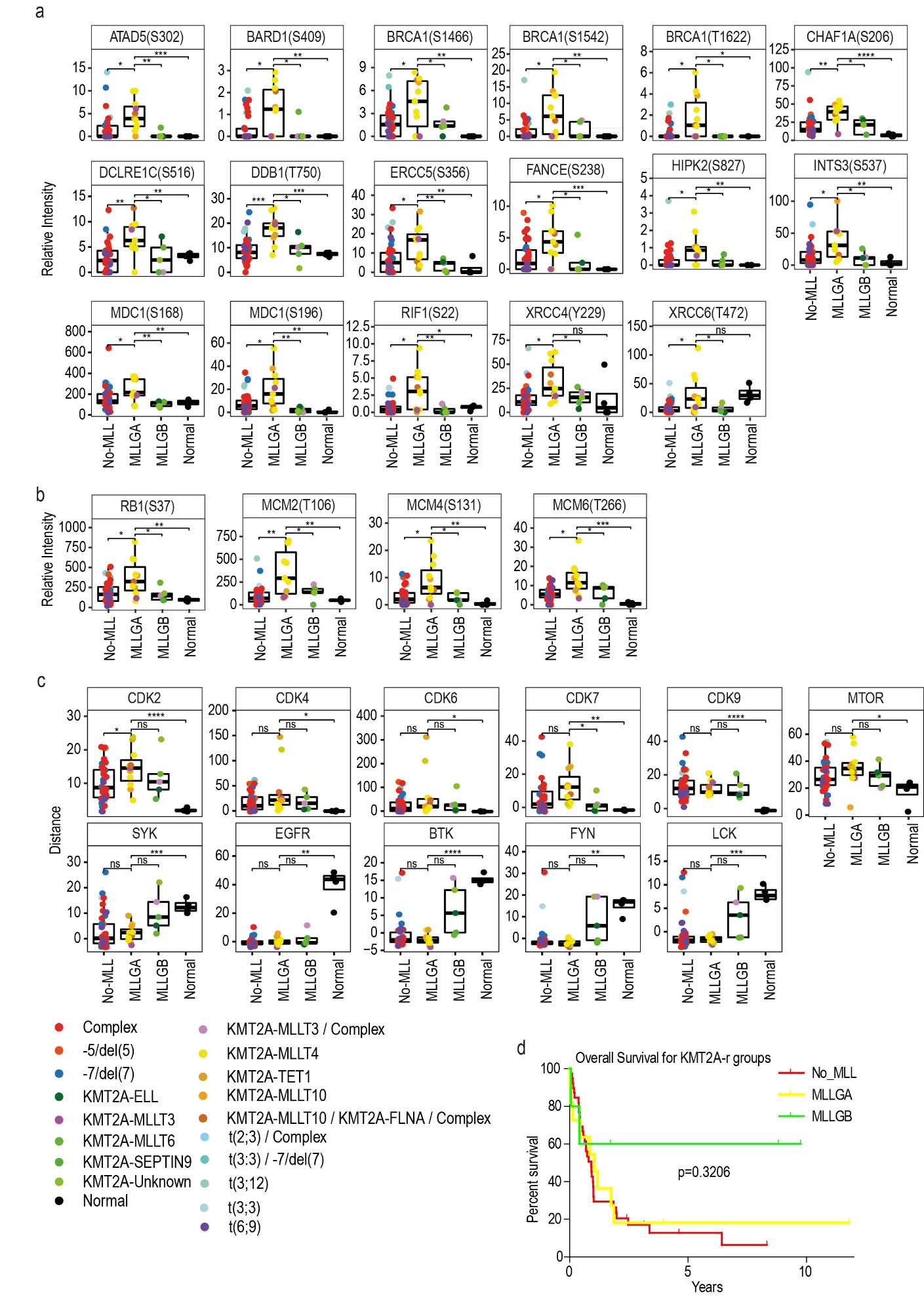

Figure. S5.

**Phosphorylation of proteins liked to DNA damage response and Replication and kinase activity as a function of KMT2A rearranged group.** **a** Phosphorylation of proteins involved in the DNA damage response as a function of KMT2A rearranged group. **b** Phosphorylation of proteins involved in replication as a function of KMT2A rearranged group. **c** KSEA estimated kinase activity a function of KMT2A rearranged group. **d** Overall survival of AML patients across KMT2A rearranged groups. Boxplots indicate median, 1^st^ and 3^rd^ quartiles. Whiskers extends from the hinge to the largest and lowest value no further than 1.5 times the distance between the 1st and 3rd quartiles. Statistical significance was calculated using unpaired two-sided Student’s t-test (**a**-**c**) or Mantle Cox test (**d**). **** p ≤ 0.0001, *** p ≤ 0.001, ** p ≤ 0.01 and * p ≤ 0.05. Normal (n=4), MLLGA (n=11), MLLGB (n=5) and No-MLL (n=39).

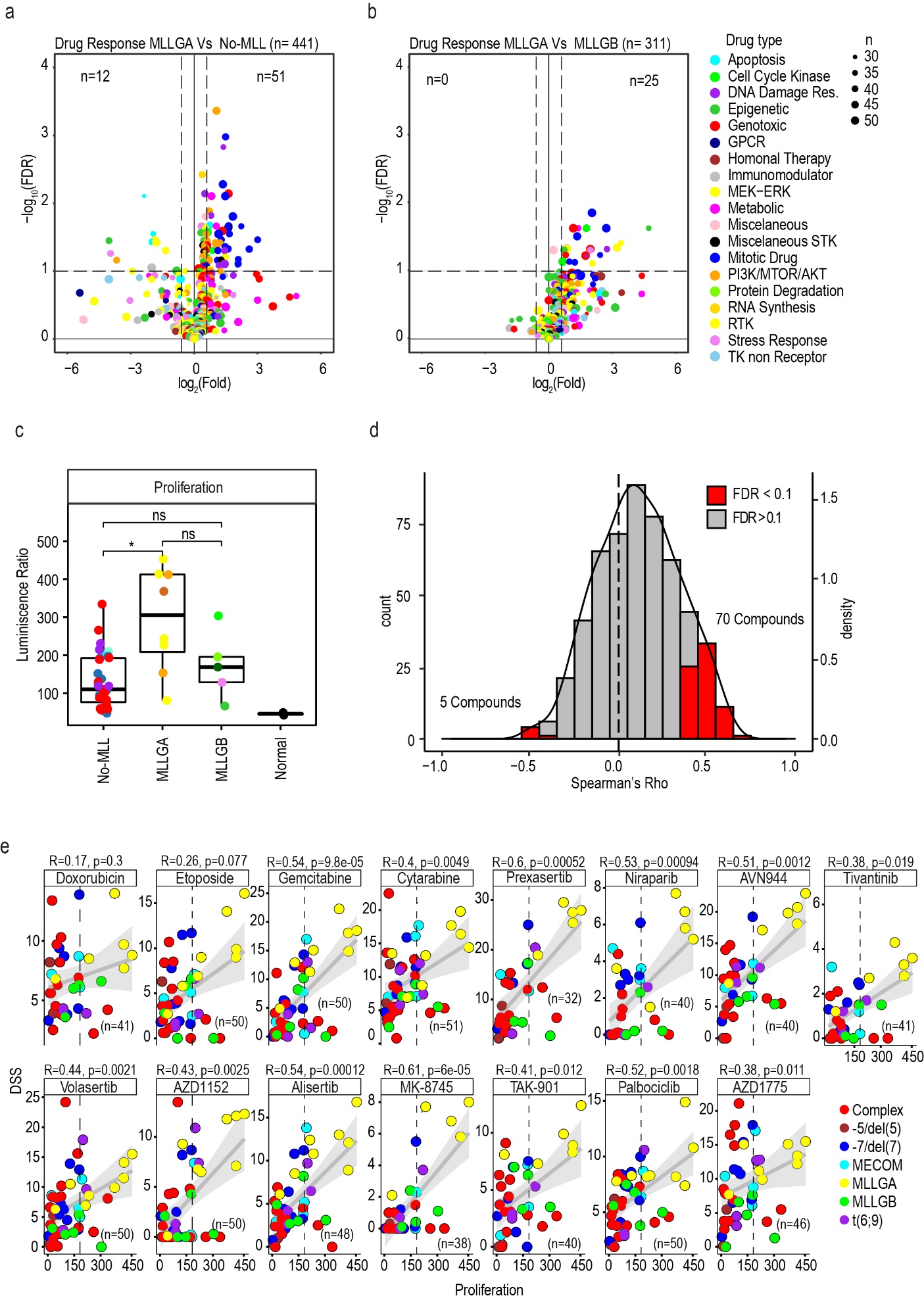

Figure. S6.

**Cellular response to compound treatment as a function of KMT2A group and correlation between proliferation and response to compound.** **a** Volcano showing compound with a significantly different response between MLLGA and No-MLL. **b** Volcano showing compound with a significantly different response between MLLGA and MLLGB. **c** Proliferation rate as a function of KMT2A rearrangement group. **d** Distribution of Spearman rank correlation values between compound response (DSS) and cell proliferation. **e** Spearman rank correlation between compound response (DSS) and proliferation for the 15 compounds that were more effective in MLLGA than in MLLGB and No-MLL. Each dot represents a compound and the size of the dot represents the number of samples used to calculate the response for that compound (**a** and **b**). Statistical significance was calculated using unpaired two-sided Student’s t-test. (**a**, **b** and **c**). FDR values were obtained by the adjustment of p-values using the Benjamini- Hochberg procedure (**a**, **b** and **d**). Each dot represents a patient sample (**c** and **e**). Boxplots indicate median, 1^st^ and 3^rd^ quartiles. Whiskers extends from the hinge to the largest and lowest value no further than 1.5 times the distance between the 1st and 3rd quartiles, *p ≤ 0.05 and Normal (n=2), MLLGA (n=8), MLLGB (n=5) and No-MLL (n=35) (**c**).

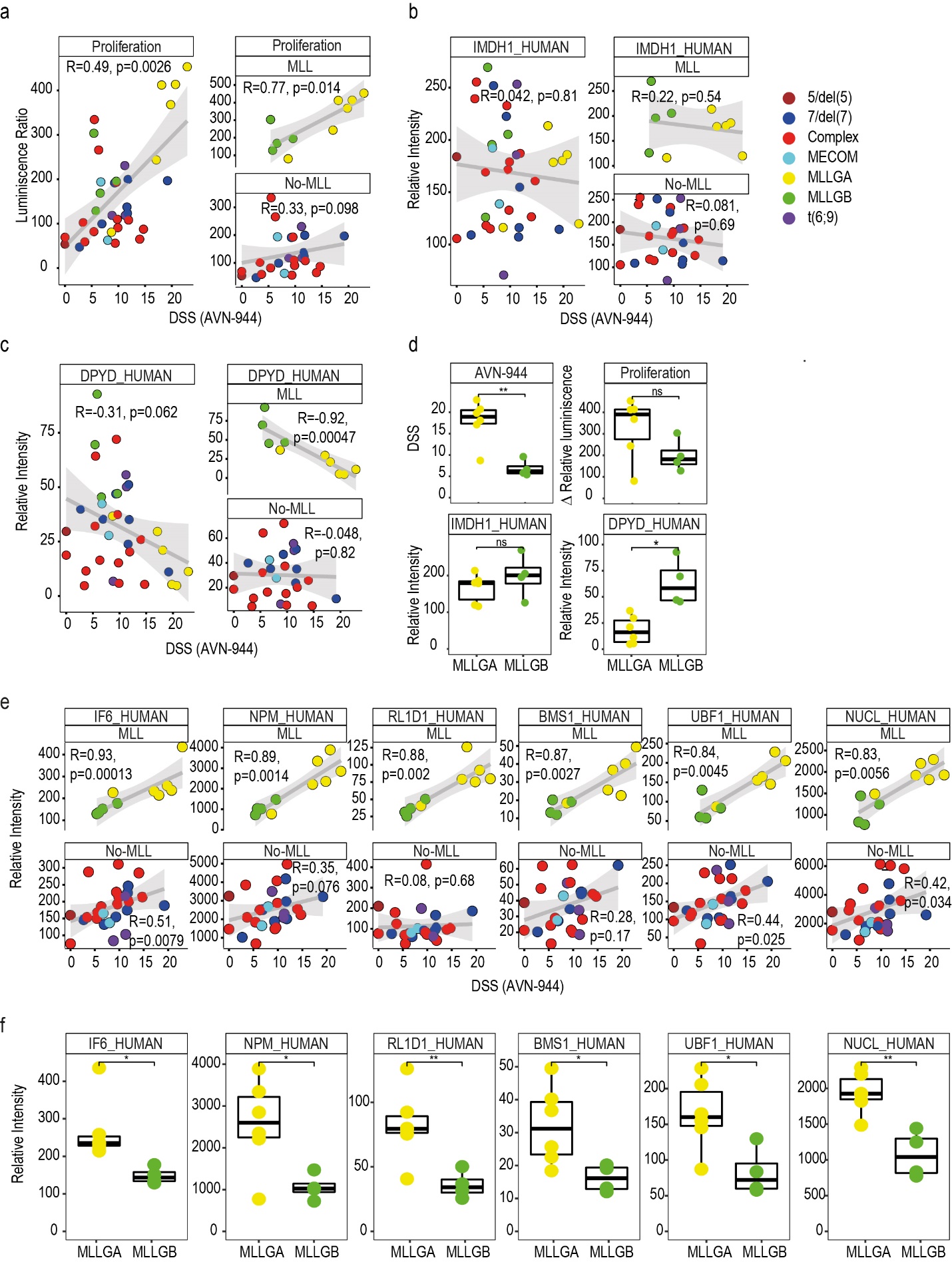

Figure. S7.

**DPYD and proteins related to the nucleolus correlate with AVN-944 DSS in KMT2A rearranged leukaemia and are differentially expressed between MLLGA and MLLGB. a** Spearman rank correlation between response to AVN-944 and proliferation. **b** Spearman rank correlation between response to AVN-944 and protein expression of IMDH1. **c** Spearman rank correlation between response to AVN-944 and protein expression of DPYD. **d** DSS for AVN-944, proliferation rate and protein expression of IMPDH1 and DPYD across MLLGA and MLLGB. **e** Spearman rank correlation between response to AVN-944 and the protein expression of the indicated genes. **f** Protein expression of nucleolar proteins. protein expression of IMPDH1 and DPYD across MLLGA and MLLGB. Boxplots indicate median, 1^st^ and 3^rd^ quartiles. Whiskers extends from the hinge to the largest and lowest value no further than 1.5 times the distance between the 1st and 3rd quartiles, statistical significance was calculated using unpaired two-sided Student’s t-test and **** p ≤ 0.0001, *** p ≤ 0.001, ** p ≤ 0.01 and * p ≤ 0.05 (**d** and **f**). MLL (n=10), No-MLL (n=26) (**a-c** and **e**). MLLGA (n=6), MLLGB (n=4) (**d** and **f**).

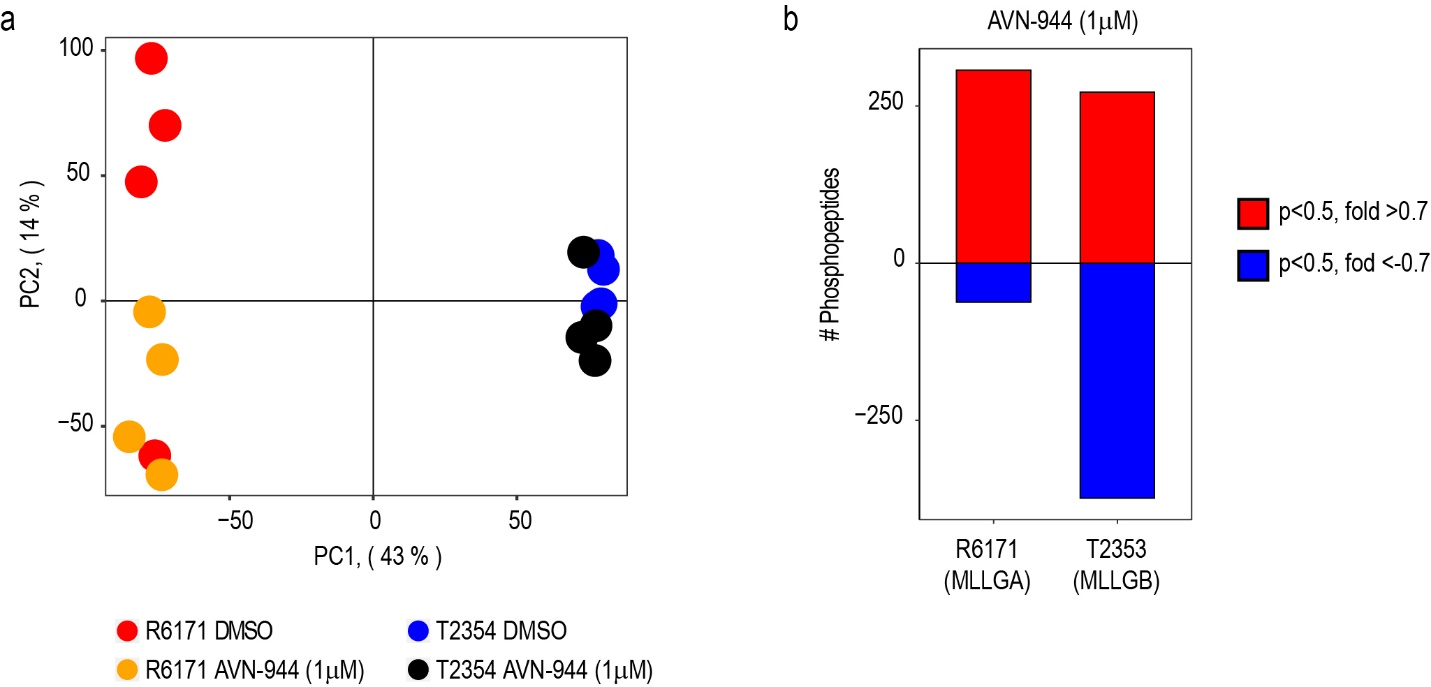

Figure. S8.

**Effect of AVN-944 treatment on the phosphoproteome of R6171 and T2354. a** PCA generated with all phosphopeptides detected after treatment of R6171 and T2354 samples with 1 µM of AVN-944 for 24 h. **b** Number of phosphopeptides significantly affected by AVN-944 treatment in R6117 and T2354 samples. Statistical significance was calculated using unpaired two-sided Student’s t-test, Phosphopeptides were counted when p-value <0.05 and fold change (log2) > 0.7 (positive values) or <-0.7 (negative values) and (n=4) independent biological replicates.
